# Promotion of Structured Motor Program Diversity Through Since-last-state Memory in *Drosophila* Larvae

**DOI:** 10.64898/2026.08.17.745218

**Authors:** William V. Smith, Stefan R. Pulver

## Abstract

Motor systems controlling locomotion must generate repetitive rhythmic activity, while also still retaining the ability to generate a diverse range of outputs. How motor systems monitor, regulate, and promote diversity of their own outputs is not well understood. Here, we perform single-step, variable-order and hidden-state Markov modelling (HSMM) on spontaneous fictive locomotor activity in the isolated *Drosophila* larval nervous system to examine how a motor system balances constraint and promotion of diversity amongst competing motor programs. We show that spontaneous fictive activity is structured by interacting mechanisms operating at multiple levels of sequence organisation. Analysis of one-step transition rules revealed a bias in activity towards activity states underlying exploration that in turn, promote transition to diverse outputs. In contrast, higher-order Markov, N-gram, and HSMM analysis indicated a memory biased towards revisiting recently executed motor programs. These mechanisms together suggest that the *Drosophila* larval locomotor system maintains a dynamic repertoire of possible motor outputs by monitoring recent activity and biasing future transitions accordingly. In this sense, fictive rhythmogenesis reflects a diversity-generating process: the larval locomotor network does not simply repeat a fixed motor programme or randomly transition from one state to another, but rather continually regulates access to rhythmic states based on recent experience. Together, these findings suggest that fictive locomotor dynamics are consistent with *adaptive* winner-takes-all competition between central pattern generating (CPG) modules that balance constraint and promotion of motor program diversity.

## Introduction

Rhythmic motor circuits balance opposing demands: they must generate stable, coordinated rhythms, while remaining close enough to a transition-ready state to flexibly access alternative motor outputs. Motor diversity is therefore not simply the production of many different behaviours, but the regulated capacity to preserve multiple possible future actions while still permitting appropriate constraint when one motor program must dominate, such as during escape or directed locomotion. This presents a central problem for rhythmogenesis: motor networks must avoid both rigid repetition and unstructured variability. How such circuits track their recent output history, regulate bias between competing motor programs, and maintain the balance between constrained state selection and flexible motor diversity remains unclear.

Across many animals, central pattern generating (CPG) networks are capable of producing different rhythmic outputs, often from the same underlying network architecture, through interactions between intrinsic circuit dynamics such as feedforward excitation and reciprocal inhibition [1], [2] and neuromodulatory inputs [3], [4]. Classic studies of the crustacean stomatogastric ganglion (STG) demonstrated that a single circuit can generate multiple motor patterns [5], [6], [7]; similar multifunctionality has been observed in invertebrate locomotor networks such as the leech crawling and swimming circuits [8] and vertebrate networks such as *Xenopus*, zebrafish, and mammalian spinal circuits [9], [10], [11], [11], [12], [13]. Rhythmic CPG circuits are generally not simply rigid oscillators but rather flexible dynamical systems capable of producing diverse coordinated outputs. However, it remains unclear how CPG circuits generate such diversity while maintaining stable coordination between a variety of mutually exclusive rhythmic outputs.

Several mechanisms have been proposed to explain how locomotor circuits transition between distinct rhythmic outputs. One possibility is that competing CPG modules interact through mutual inhibition, enabling action selection between antagonistic motor programs [7]. Alternatively, slow adaptive processes or intrinsic memory mechanisms influence the probability of future motor states. Activity-dependent processes such as Na⁺/K⁺ pump–mediated afterhyperpolarisation can generate short-term motor memory in rhythmic circuits [14], [15], [16], while neuromodulators can reconfigure network excitability and alter locomotor dynamics [17]. More broadly, locomotor circuits often contain interneuron populations selectively recruited into distinct oscillatory modules responsible for different coordination patterns or speeds [18], [19], [20], [21]. Despite these insights, principles underlying how locomotor circuits generate diverse rhythmic outputs while maintaining reliable transitions between them remains elusive. In particular, it remains unclear how intrinsic circuit dynamics regulate transitions amongst multiple possible rhythms over long time courses. Additionally, it remains unclear how short-term behavioural history or transitions between competing motor programs are regulated to maintain optimal motor output.

Locomotion involves exclusion of competing motor output likely via ‘winner-takes-all’ mechanisms, where one motor program is selected and others supressed to ensure execution of appropriate locomotor behaviours at a given time. Appropriate rhythmogenesis involves an impeded orchestration of the motor system to create a smooth and necessitate motor output. However, how the motor system balances the need for appropriate rhythmogenesis with potentiation of diverse motor outputs is complex. Conceptually, command-like neuron structures may be in persistent winner-takes-all competition to generate exclusive motor output [22], [23]. Winner-take-all models within action selection embed mutual- and self-inhibitory dynamics to ensure dynamic switching in outputs [24], [25], use asymmetric coupling or gain-modulation between command-like structures to generate stochastic or external-driven structure to motor output [26], and can demonstrate history-dependent modifications to output based on neuromodulatory gain, plasticity, or fatigue dynamics [27], [28], [29], [30]. Here, we utilise the *Drosophila* fictive motor system to examine whether motor systems contain such proposed intrinsic mechanisms to promote structured access to diverse motor states in line with winner-takes-all competitive frameworks.

Here, we investigate the intrinsic functional organisation of locomotor dynamics in the isolated *Drosophila* larval nervous system to examine the hierarchical organisation of rhythmogenesis. We show that spontaneous fictive motor sequences are non-random and exhibit higher-order transition structure governed by short-term recurrence instance-since-last-state memory. Furthermore, fictive locomotor activity occupies distinct coordination regimes and contains identifiable hidden macrostates that bias transition probabilities between motor programs. Within this organisation, anterior asymmetric-rich states – reflective of larval exploratory headsweeps – occupy hub-like positions in the transition landscape, suggesting that asymmetric activity may organise access to alternative fictive trajectory, thus promoting output diversity. Together, these findings suggest that fictive locomotor dynamics may emerge from *adaptive* winner-takes-all competition between CPG modules that simultaneously generate and constrain motor diversity. From these results, we propose a set of testable predictions for adaptive winner-takes-all competition between CPG modules and identify connectome-constrained candidate neurons that may organise and modulate motor competition in the *Drosophila* larval locomotor system. Overall, in the absence of sensory feedback and descending goal-oriented modulation, spontaneous fictive output demonstrates intrinsic tendencies of the motor network to generate, revisit, and reorganise motor patterns towards output diversity.

## Results

### Fictive Motor Activity is Non-Random and Organised Around Distal Transition Hubs

The isolated *Drosophila* ventral nerve cord (VNC) spontaneously generates rhythmic activity representative of intact behaviours which can be measured by expressing genetically-encoded calcium indicators in glutamatergic neurons [31], [32]. Previous work has characterised the reliable correspondence between activity patterns of the isolated *Drosophila* CNS, i.e., “fictive” activity patterns, and intact animal behaviour [1], [31], [33]. We categorised neural activity in the VNC to define fictive motor programs: distal activities, i.e., thoracic fictive asymmetries (AS), thoracic anterior symmetric bursts (AB), and abdominal symmetric posterior bursts (PB), and wave activities, i.e., fictive forward (FW) and backward (BW) waves (Figure 1A). Previously, our work [1] demonstrated and quantified the diversity and variability in motor output in the isolated VNC, and showed that motor programs typically do not overlap with one another in time under normal physiological conditions. However, the dynamical principles governing how this diversity is generated and maintained remain unclear. To understand if the isolated motor network demonstrates consistent dynamical rules, we first evaluated whether there are reproducible relationships and transition patterns between fictive motor programs, or if fictive output is simply stochastic.

**Figure 1.**
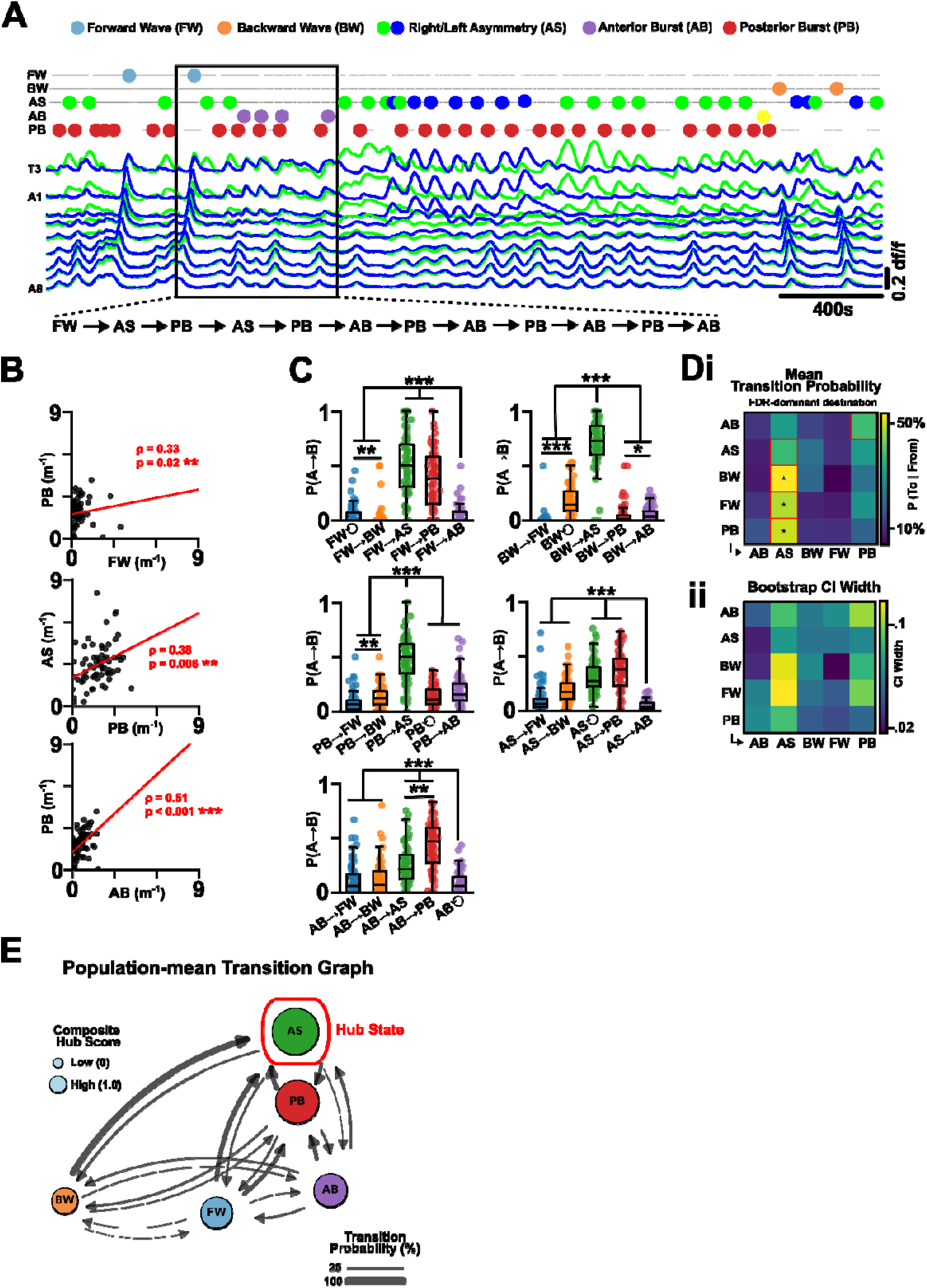
Spontaneous Fictive Dynamics Are Non-Random, Oriented Towards High Entropic Distal Activity Patterns. (**A**) A representative calcium imaging trace of CNS activity from an isolated preparation (total N = 69, 45 minutes) with annotated sequences of fictive activity patterns (Forward = FW, Backward = BW, Anterior Asymmetric = AS, Anterior Symmetric = AB, Posterior Burst = PB). Coloured dots indicate incidence of each fictive behaviour. Box provides a representative example the fictive dynamic sequence generated from the calcium imaging data. (**B**) Significant proportionality between frequency (m) of fictive activity patterns (only significant correlations are shown). (**C**) Transition probabilities between each fictive state separated into each preceding fictive state (e.g., FW → X). (**Di**) The mean transition probability between fictive states taken from average of transition dynamics within each isolated preparation alongside (**Dii**) bootstrapped confidence intervals for each transition. (**E**) Population-mean transition graph between fictive states whereby transition probabilities are presented as averages of individual preparation averages. Node size represents the composite hub score – the z-scored and averaged inflow and outflow dynamic properties of the state – with arrow thickness designating transition probability. Nodes are organised along the x-axis by increasing outflow entropy. Note that distal activity states – AS/PB – represent dominant convergent and throughput states within fictive network dynamics. Post-hoc p-values: * p < .05, ** p < .01, *** p < .001

To explore the fictive system’s spontaneous transition dynamics, we imaged and scored fictive locomotor activity patterns in 3rd instar *Drosophila* larvae (N = 69) to generate sequence series of fictive behaviours for each preparation (Figure 1A). As we reported [1], there is considerable diversity and variability within the output of the fictive motor system. Across preparations, AS was the most frequently observed fictive activity pattern, followed by PB, BW, AB, and FW (AS = 2.96 ± 0.18 min⁻¹; PB = 1.84 ± 0.12 min⁻¹; BW = 0.99 ± 0.07 min⁻¹; AB = 0.62 ± 0.06 min⁻¹; FW = 0.53 ± 0.07 min⁻¹; Figure 1B). Analysis of fictive behavioural frequencies revealed that relationships between fictive motor programs were selective rather than global. In particular, PB frequency was positively correlated with the frequency of AB (ρ = 0.51, q < .001), AS (ρ = 0.38, q = 0.006), and FW (ρ = 0.33, q = 0.019) (Figure 1C; Supplementary Table 1). In contrast, BW frequency did not significantly correlate with other fictive behaviours, and other relationships, including FW-AS, FW-AB, BW-PB, and AB-AS, were not significant after correction (Supplementary Table 1). Thus, the proportional relationships between fictive behaviours were not simply explained by global variation in activity across different preparations. Instead, specific fictive behaviours covaried with one another, suggesting that spontaneous rhythmogenesis occupies a structured, but variable, behavioural landscape.

Given that specific fictive motor programs covary in frequency, we next evaluated the sequential organisation of fictive behaviours through transition analysis. Transition probabilities were computed independently for each preparation and then averaged across the population. Fictive transitions were strongly non-uniform, with each behavioural class showing a preferred or high-probability successor state (Figure 1D; Supplementary Figure 2, 3). FW most commonly transitioned into AS (50.7 ± 3.3%), with PB as a strong secondary target (36.8 ± 3.3%). BW showed the most selective transition profile, transitioning predominantly into AS (70.6 ± 2.6%), with comparatively weaker BW self-transitioning (17.9 ± 1.7%). PB also preferentially transitioned into AS (47.7 ± 2.4%), while AB most commonly transitioned into PB (42.1 ± 2.7%). AS, in turn, most frequently transitioned into PB (36.1 ± 2.1%) and also showed substantial self-transitioning (30.9 ± 1.8%). Overall, transition probabilities were not uniformly distributed across successor states, demonstrating that spontaneous fictive activity evolves through reproducible and non-random transition rules.

To move beyond one-step fictive transition preferences, we next analysed the fictive transition matrix as a directed weighted graph, with fictive motor programs represented as nodes and transition probabilities represented as edges. We explored whether particular fictive states occupy hub-like positions within the global transition landscape. By examining a variety of transition metrics, including incoming transition strength, outgoing transition structure, and bridge-like centrality, we produced a composite hub score to determine which fictive states combine strong connectivity with central positions in the transition network (Figure 1E). This analysis indicated that fictive dynamics are not organised as a simple linear chain of motor programs. Instead, the transition network is structured around distal activity states, particularly AS and PB. AS showed especially prominent hub-like properties, consistent with its role as a convergence state for FW, BW, and PB transitions, while also redistributing activity into PB or back into AS. PB also occupied a high-connectivity position, receiving input from FW, AS, and AB, and then transitioning strongly into AS. Thus, AS and PB together define a distal hub-like subspace through which spontaneous fictive activity is repeatedly routed.

Overall, spontaneous fictive motor activity in the isolated VNC is diverse but not random. The fictive network exhibits reproducible frequency relationships and transition rules, with distal activity patterns – especially AS and PB – acting as central organising states in the fictive motor landscape.

### Fictive Locomotory Dynamics Contain Structured, Multi-step Transition Sequences Not Captured by First-order Markov Dynamics

To assess the predictive utility of consistent one-step transition dynamics, we quantified the occurrence of multi-step fictive motifs and compared their empirical frequency to expectations derived from first-order Markov dynamics (Figure 2). First, we evaluated the predictive accuracy of first-order transitions on longer fictive behavioural motifs. For increasing N-gram length, the prediction error between empirical motif frequencies and those expected from one-step transition rules increased systematically with motif length, with summed absolute prediction error increasing from ∼0 for 2-grams to 0.28 for 3-grams, 0.46 for 4-grams, 0.67 for 7-grams, and 0.98 for 12-grams (Figure 2Ai). Consistently, the correlation between empirical and Markov-predicted motif probabilities declined as motif length increased, from r = 1.00 for 2-grams, to r = 0.95 for 3-grams, r = 0.88 for 4-grams, r = 0.60 for 7-grams, and r = 0.21 for 12-grams (Figure 2Aii). Thus, a one-step transition rule reliably captures the immediate short motif structure of the fictive system, but its predictive accuracy progressively weakens for longer fictive trajectories. These data indicate that fictive behavioural sequences are not fully described by the immediately preceding state alone.

**Figure 2.**
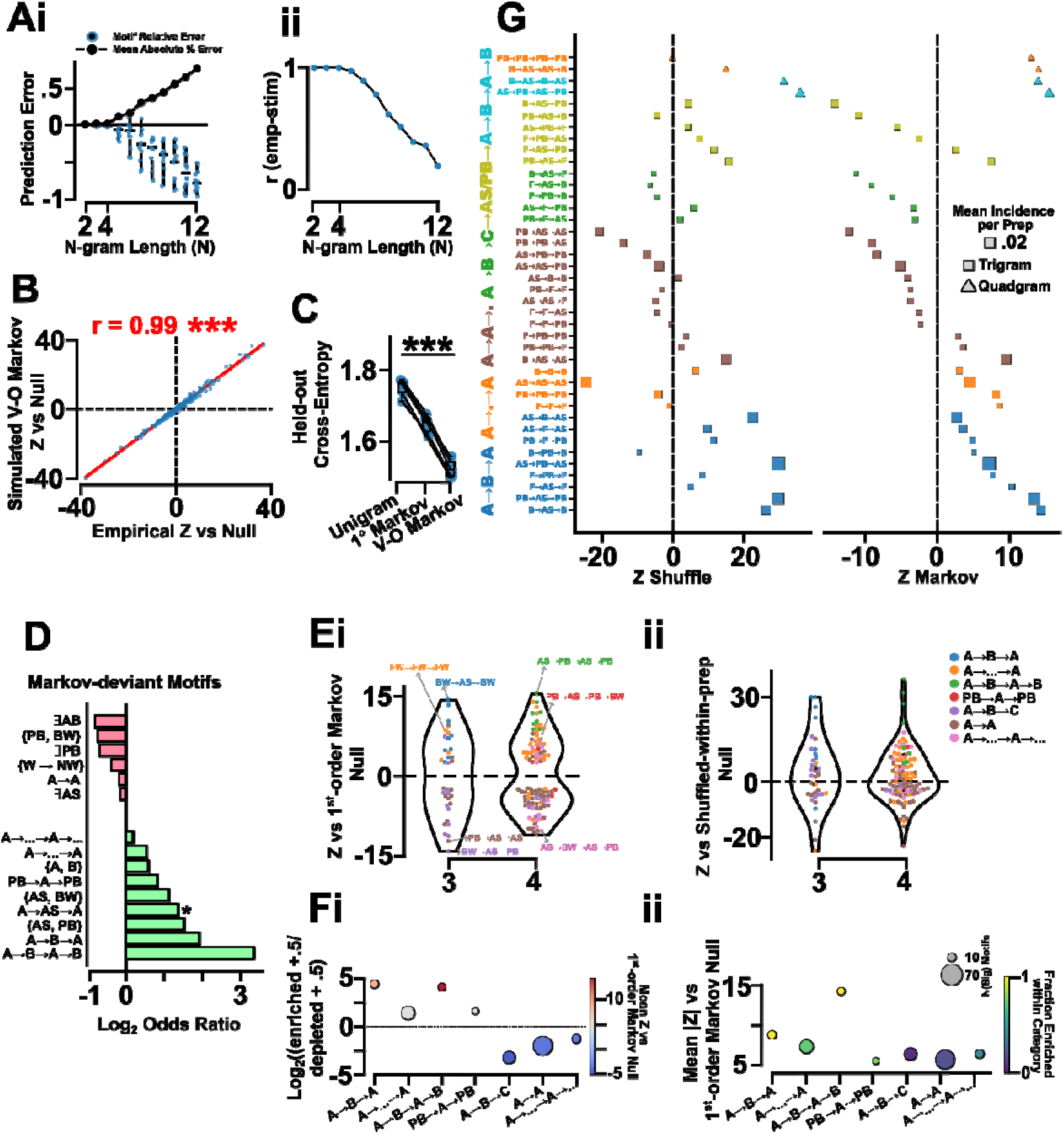
Multi-step Fictive Motifs Occur Beyond Predictions from one-step Transition Probabilities. (**A**) The prediction error (**i**) and Pearson’s correlation coefficient (**ii**) of a first-order Markov model for lengths of fictive sequences (n-gram length 2-12) showing a deviation in predictive power beyond n-gram length of 4. (**B**) The correspondence between all present empirical multi-step motifs and variable-order Markov model (maximum order = 3 and backoff) simulated multi-step motifs compared against the mean count of said motif across shuffled sequences where each preparation’s fictive behavioural order is randomly permuted. (**C**) Hold-out cross-entropy of unigram (i.e., transition dependent on global fictive frequencies), first-order Markov model (i.e., transition dependent on prior fictive state), and variable-order Markov model (i.e. transition dependent on the history of 3 prior fictive states). Held-out cross-entropy captures the ability of a probabilistic model to predict unseen, non-training data (5 equal groups). Low cross-entropies indicate models with better predictions. (**D**) Multi-step fictive motifs whose occurrence deviates from expected appearance predicted by a first-order Markov model. Three categorical motif types are underrepresented relative to a first-order Markov prediction: contains an anterior burst in the sequence (AB), has a repeating fictive event anywhere (A→…→A→…), has an adjacent repeating fictive event (A→A). Thirteen categorical motif types are overrepresented relative to a first-order Markov prediction: contains an anterior asymmetric event (lllAS), a chain of different fictive events (A→B→C), contains a posterior burst event (PB), fictive motif where the 1 fictive event is the same as the final fictive event (A→…→A), motifs exclusively involving posterior burst and fictive backward events ({PB, BW}), motifs exclusively alternate between fictive wave and distal fictive events ({W, NW}), motifs that start and end with a posterior burst (PB→…→PB), motifs with an anterior asymmetric event contained in the motif (A→AS→A), motifs exclusively involving anterior asymmetric events and fictive backward events ({AS, BW}), motifs containing exclusively only two different fictive events ({A,B}), motifs exclusively involving anterior asymmetric events and posterior bursts ({AS, PB}), motifs that alternative between two different fictive events (A→B→A→B), three-step motif where the a type of fictive event starts the motif and that state reappears after an intermittent fictive event (A→B→A). (**Ei**) The distribution of motif enrichment (Z-scores) relative to the first-order Markov null for tri-gram (N = 3) and quad-gram (N = 4) motifs. Positive values indicate motifs occurring more frequently than predicted by pairwise transition probabilities, whereas negative values indicate motifs occurring less frequently than expected. Points represent individual motifs coloured by motif class. (**Eii**) Motif enrichment relative to a shuffle-within-preparation null preserves behavioural composition but randomises order. Z-scores indicate deviations from random behavioural ordering, confirming structured sequential organisation in fictive behaviour. (**Fi**) Enrichment magnitude for representative motif classes. Points show log₂ enrichment relative to the null model, coloured by mean Z-score relative to the first-order Markov null. (**Fii**) Mean absolute enrichment (|Z|) relative to the first-order Markov null across motif classes. Point size indicates the number of motifs per class, and colour indicates the fraction of motifs significantly enriched relative to the shuffle null.

To reinforce the predictive utility of wider sequence context, we compared the predictive performance of models with different levels of sequential information using held-out cross-entropy (Figure 2B,C). A unigram model, which only preserves global fictive behavioural frequencies, performed worst across held-out folds (mean cross-entropy = 1.75). Incorporating the immediately preceding fictive state improved prediction under a first-order Markov model (mean cross-entropy = 1.65), while a variable-order Markov model incorporating longer sequential context performed best numerically (mean cross-entropy = 1.52). Thus, including multi-step transition context reduced mean predictive error by 0.23 relative to the unigram model and by 0.12 relative to the first-order Markov model.

To explore the multi-step fictive motif structure directly, we systematically identified motifs whose empirical occurrence deviated from first-order Markov expectations (Figure 2D). As expected, 2-gram motifs did not significantly deviate from the bigram null, since these are the transition probabilities directly preserved by the model. In contrast, higher-order motifs showed substantial deviation from first-order expectations. For 3-grams, 36 motifs were significantly Markov-deviant after FDR correction, with an equal split between enriched and depleted motifs (18 enriched, 18 depleted). For 4-grams, 111 motifs were significantly deviant, comprising 51 enriched and 60 depleted motifs. Thus, higher-order deviations were not a global inflation of all longer sequences, but instead reflected a selective structure in which particular fictive trajectories occurred more or less often than expected from one-step transition probabilities alone.

Several enriched motifs were centred on repeated returns through distal activity states. For example, BW→AS→BW was strongly enriched relative to the first-order null (Z = 14.37, q = 0.005), as were PB→AS→PB (Z = 13.42, q = 0.005) and FW→AS→FW (Z = 10.34, q = 0.005). This structure was also prominent in 4-gram motifs, with AS→PB→AS→PB (Z = 15.50, q = 0.005), BW→AS→BW→AS (Z = 13.96, q = 0.005), and PB→AS→PB→AS (Z = 11.84, q = 0.005) occurring considerably more often than predicted by first-order Markov dynamics. Conversely, several motifs were significantly depleted, including BW→AS→PB (Z = −14.13, q = 0.002), PB→AS→AS (Z = −12.19, q = 0.005), PB→AS→BW (Z = −10.88, q = 0.002), and PB→AS→AS→AS (Z = −11.04, q = 0.005). Therefore, AS does not act simply as a non-specific intermediate state. Instead, the behavioural state preceding AS shapes which fictive trajectory is likely to follow, indicating that the fictive network retains information about prior behavioural context.

To further examine the structure of these deviations, we analysed motif enrichment across sequence categories (Figure 2E,F). Alternating ABA-like tri-grams were uniformly enriched, with all 9 significant motifs in this class showing positive enrichment (median Z = 7.20). Similarly, ABAB-like oscillatory motifs were also consistently enriched, with all 8 significant motifs showing positive enrichment and a high median Markov-deviation score (median Z = 11.09). Return-to-state motifs were predominantly enriched, with 26 of 31 significant motifs showing positive enrichment. In contrast, progression-chain motifs were predominantly depleted, with 14 of 16 significant motifs occurring less often than expected (median Z = −5.07), and adjacent-repeat chains were also mainly depleted, with 42 of 52 significant motifs showing negative enrichment (median Z = −3.68). Thus, the higher-order structure of fictive activity is not simply an increased tendency to repeat behaviours, but instead reflects selective recurrence, alternation, and return-to-state dynamics within particular fictive trajectories.

Importantly, we next tested whether these Markov-deviant motifs were simply an artefact of averaging across preparations or of slow changes in behavioural composition over time (Supplementary Figure 2). Under the original global stationary Markov null, 52 tri-gram motifs were significantly deviant, comprising 27 enriched and 25 suppressed motifs, while 164 quad-gram motifs were significantly deviant, comprising 89 enriched and 75 suppressed motifs. Comparison with a population-level non-stationary Markov null produced almost identical enrichment structure: 51 of 52 significant tri-grams were retained (98.1% retention), and 156 of 164 significant quad-grams were retained (95.1% retention). Enrichment scores were also almost perfectly correlated between the original and population-level non-stationary nulls for both tri-grams (ρ = 0.99) and quad-grams (ρ = 0.99). McNemar tests indicated no systematic shift in significance classification for either motif length. Thus, the majority of Markov-deviant motif structure cannot be explained simply by population-level non-stationarity in behavioural state usage.

We then applied stricter preparation-specific Markov nulls, which account for the fact that individual preparations can have different transition biases. As expected, these stricter nulls reduced the number of retained significant motifs, especially for 4-grams. Under the preparation-specific stationary null, 34 of 52 tri-gram motifs were retained (65.4% retention), and 92 of 164 quad-gram motifs were retained (56.1% retention). Enrichment scores nevertheless remained strongly correlated with the original global Markov null for both 3-grams (ρ = 0.80) and 4-grams (ρ = 0.88). Under the strictest preparation-specific non-stationary null, 32 of 52 tri-grams were retained (61.5% retention), and 78 of 164 quad-grams were retained (47.6% retention). Even here, enrichment scores remained positively correlated with the original global null for both 3-grams (ρ = 0.6) and 4-grams (ρ = 0.79). Therefore, preparation-specific transition structure and temporal drift explain part of the original motif enrichment landscape, particularly among longer 4-gram sequences, but they do not eliminate higher-order structure. Instead, they separate a broad population-level motif structure from a more conservative core of motifs that remain Markov-deviant even after accounting for each preparation’s own transition dynamics.

To ensure that these motif structures were not driven by a small number of preparations, we also evaluated motif stability using a leave-one-preparation-out analysis (Figure 2Fii). The majority of motifs showed stable enrichment structure across preparations, with 247 of 299 motifs classified as highly stable. This stability was especially strong for shorter motifs, with 16 of 16 2-grams and 59 of 63 3-grams classified as highly stable, and remained substantial for 4-grams, where 172 of 220 motifs were stable. Importantly, the strongest enriched motifs, including AS→PB→AS→PB, BW→AS→BW→AS, PB→AS→PB, and AS→PB→AS, retained their enrichment direction across leave-one-preparation-out iterations. Therefore, these higher-order motif structures are not explained by a single dominant preparation, but instead reflect reproducible features of the fictive locomotor circuit.

The largest Markov-deviant motif structures are shown in Figure 2G. Together, these analyses indicate that fictive motor dynamics are not fully captured by either random fictive behavioural occurrence or first-order Markov transition rules. Instead, specific multi-step fictive motifs are enriched or suppressed relative to first-order expectations, indicating that the current state of the fictive network retains information about prior fictive behavioural context.

### Fictive Dynamics are Non-Stationary with Critical Change-points Oriented Towards Distal Activities

The isolated fictive motor network may exhibit slow changes in fictive behavioural organisation over time due to processes such as neuronal fatigue, adaptation, or underlying state changes. Before interpreting fictive sequence dynamics mechanistically, it was important to determine whether fictive activity remains stable over time or whether any drift is a product of activity decay. To address this, we performed stationarity analysis to evaluate whether fictive behavioural structure remains constant across recordings.

Stationarity analysis demonstrated that the isolated fictive motor network exhibits moderate drift in fictive behavioural structure that is inconsistent with simple neural fatigue or degenerative decay, but instead reflects a dynamically evolving system. Comparing the transition structure between the 1^st^ and 2^nd^ halves of each recording revealed that nearly all preparations exhibited some level of divergence in fictive transition probabilities across time (median JS divergence = 0.34 ± 0.01; N = 69) (Figure 3A). Thus, even over coarse temporal partitions, fictive activity did not remain fully stationary. To examine this drift more closely, we analysed behavioural transitions using sliding windows across each sequence. Across the population, most preparations showed moderate continuous drift in transition structure (median JS divergence = 0.31 ± 0.01) (Figure 3Bi), with the median level of drift reproducible across preparations (Figure 3Bii). Importantly, drift magnitude was independent of sequence length (r = 0.10, p = 0.422) (Figure 3Biii), indicating that the observed divergence does not arise simply from longer recordings accumulating greater variability. Instead, fictive behavioural organisation evolves intermittently across time within preparations (Figure 3Biv), suggesting that the system does not operate under a stationary Markov process with fixed transition probabilities. Consistent with this, 82.1% of preparations were classified as showing moderate drift, while only 11.9% were classified as stationary and 6.0% as strong-drift preparations (Figure 3Ci).

**Figure 3.**
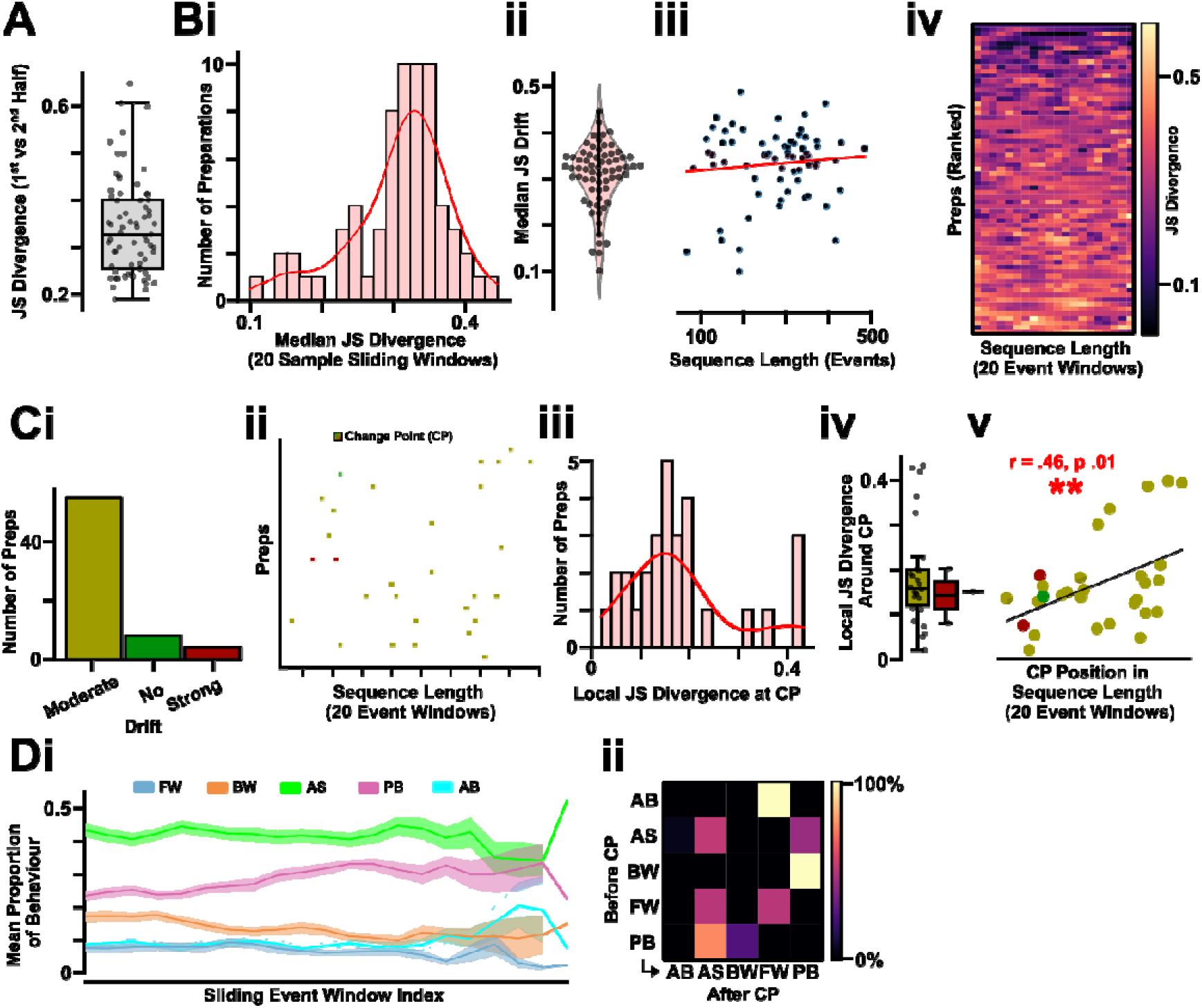
Fictive Activity Moderately Drifts Over Recording Time with Change Points Orientating the Fictive Network to Distal Activity Patterns. (**A**) Jensen–Shannon (JS) divergence of the behavioural transition distribution between the 1^st^ and 2^nd^ half of each isolated preparation recording period. (**B**) The median JS divergence between the behavioural transition distribution (**i**) quantified across preparations within 20 sample sliding windows, (**ii**) overall in the population dataset, and (**iii**) across sequence length. (**Biv**) The median JS divergence within each 20-sample sliding window of all ranked preparations across normalised time. (**C**) The quantity of isolated preparation demonstrating no, moderate, or strong drift in observed fictive behaviours. (**Cii**) The statistically-significant change point/s (CP) exhibited by preparations (N=36/69) across time normalised to the sequence position. (**Ciii**) The local JS divergence between the observe behaviour probability transition distribution before vs after the CP. (**Civ**) The local JS divergence between the observed behaviour probability transition distribution of preparations classified as exhibiting no, moderate, and strong drift. (**Cv**) The relationship between the local JS divergence around the CP compared against the normalised temporal positioning of the CP in the recording period. (**Di**) The changing mean proportion of fictive behaviours (FW = forward, BW = backward wave, AS = anterior asymmetry, PB = posterior burst, AB = anterior symmetric burst) across the time of each recording normalised by sliding windows. (**Cii**) A transition probability between the fictive behaviour before and after a statistically-significant critical transition point across the population dataset where CP/s exist.

While gradual drift characterised the majority of preparations, a subset of recordings exhibited more abrupt reconfigurations of fictive behavioural structure, detectable as discrete change points (Figure 3Cii). In total, 29 change points were detected across 18 preparations. These local reorganisations were generally moderate in magnitude (median local JS divergence = 0.18 ± 0.02), although some events reflected larger changes in the local fictive landscape (maximum local JS divergence = 0.43) (Figure 3Ciii). Notably, the magnitude of these local reorganisations was not significantly different across preparation-level drift classes (Figure 3Civ). However, local JS divergence increased later in the sequence (r = 0.46, p = 0.012), indicating that larger local reorganisations were more likely to occur as recordings progressed (Figure 3Cv). Independent of these discrete change points, we also observed gradual shifts in the mean proportions of fictive behaviours across the recording period (Figure 3Di). When abrupt reorganisations did occur, they were strongly centred around distal activity regimes. The most common dominant-behaviour patterns at change points were AS→AS (11/29, 37.9%), AS→PB (9/29, 31.0%), and PB→AS (3/29, 10.3%) (Figure 3Dii). Thus, even when local reorganisation occurred, it did not reflect a random redistribution across all fictive motor programs. Instead, abrupt changes in the fictive landscape largely reflected persistence within, or redistribution between, AS and PB-dominated regimes.

The non-stationary nature of fictive dynamics could, in principle, generate the multi-step motif features identified in the original population-level analysis. To evaluate this, we compared motif enrichment structure under the original stationary Markov null with population- and preparation-level non-stationary Markov controls (Supplementary Figure 2). Under the original global stationary Markov null, 52 tri-gram motifs were significantly deviant, comprising 27 enriched and 25 suppressed motifs, while 164 quad-gram motifs were significantly deviant, comprising 89 enriched and 75 suppressed motifs. Population-level non-stationary controls had minimal effect on this motif landscape, preserving 51/52 significant 3-grams (98.1%) and 156/164 significant 4-grams (95.1%) identified by the original analysis. Consistently, motif enrichment Z-scores showed near-perfect concordance between the original stationary and population-level non-stationary analyses for both 3-grams (ρ = 0.998) and 4-grams (ρ = 0.999), with no retained motifs switching enrichment direction. McNemar tests also indicated no systematic shift in significance classification for either motif length. Thus, population-level temporal drift does not explain the higher-order motif structure.

In contrast, preparation-specific controls substantially reshaped the longer motif landscape. Under a preparation-specific stationary Markov null, 34/52 original significant 3-grams were retained (65.4%) and 92/164 original significant 4-grams were retained (56.1%), with enrichment scores remaining strongly correlated with the original global null for both 3-grams (ρ = 0.799) and 4-grams (ρ = 0.884). Under the strictest preparation-specific non-stationary null, 32/52 original significant 3-grams were retained (61.5%), with 20 motifs lost and 35 newly significant motifs detected. For 4-grams, 78/164 original significant motifs were retained (47.6%), with 86 motifs lost and 33 newly significant motifs detected. This represented a significant shift in 4-gram motif-significance structure (McNemar p = 1.0 × 10⁻⁶), while the shift for 3-grams did not reach significance (p = 0.058). Therefore, some longer motifs detected under the original global Markov null reflect preparation-specific transition structure and time-varying first-order regimes. Nonetheless, a substantial subset of motifs survived this strictest evaluation, with 31/32 retained 3-grams and 77/78 retained 4-grams preserving the same enrichment direction. These retained motifs therefore represent strong evidence for higher-order sequence structure that cannot be explained by non-stationary first-order dynamics alone.

Taken together, fictive motor output does not appear governed by a stationary transition process, but instead evolves through a combination of gradual drift and intermittent state reorganisation. Given that drift magnitude was not explained by sequence length, and that fictive behavioural changes were directed toward specific distal activity regimes rather than broad loss of activity, non-stationarity is unlikely to reflect circuit fatigue alone. Rather, the fictive network appears to evolve through distinct dynamical modes centred around AS/PB fictive activity patterns.

### Categorical Since-Last-State Memory Predicts Multi-Step Fictive Motif Structure

The preceding analyses demonstrated that fictive motor dynamics contain non-stationary, multi-step motifs not captured by first-order Markov model dynamics. In effect, the probability of a particular future fictive state occurring depends upon recent fictive behavioural history. Such history-dependent modulation of activity is not solely predictable by knowing the immediately preceding state or global fictive frequencies. To identify the type of memory structure, we next explored a series of logistic-regression models incorporating different forms of fictive state historical dependency.

We compared logistic-regression models encoding distinct forms of fictive behavioural history, including previous state identity, persistence, state-specific recency, and recent sequence diversity. Across 18,338 held-out transition events from 69 preparations, fictive evolution was best explained by a single interpretable memory variable: the number of intervening fictive events since each state last occurred in the sequence (Figure 4A). We considered six logistic-regression transition models, each corresponding to distinct mechanistic hypotheses about the putative underlying network dynamics. The full memory model, which combined multiple forms of recent history, performed best, producing the lowest held-out log loss (M6 = 1.195) and highest accuracy (0.523). However, the since-last-state model performed nearly as well while using a substantially simpler parameterisation. Model 4 reduced held-out log loss from 1.305 under the current-state model to 1.203, increased accuracy from 0.460 to 0.516, and explained 7.76% deviance relative to the no-memory current-state model (Figure 4Bi). By comparison, the full memory model explained 8.43% deviance. Thus, the since-last-state model captured approximately 92% of the predictive gain achieved by the full memory model, despite using less than half the approximate number of independent parameters (M4 = 44; M6 = 96). Similarly, model 4 reproduced Markov-deviant motif structure (ρ = 0.935), close to the full memory model (ρ = 0.949), and higher than the current-state model alone (ρ = 0.773) (Figure 4Bii). Consequently, since-last-state memory represents the best interpretable single-memory model for capturing the statistical structure of fictive behavioural sequences.

**Figure 4.**
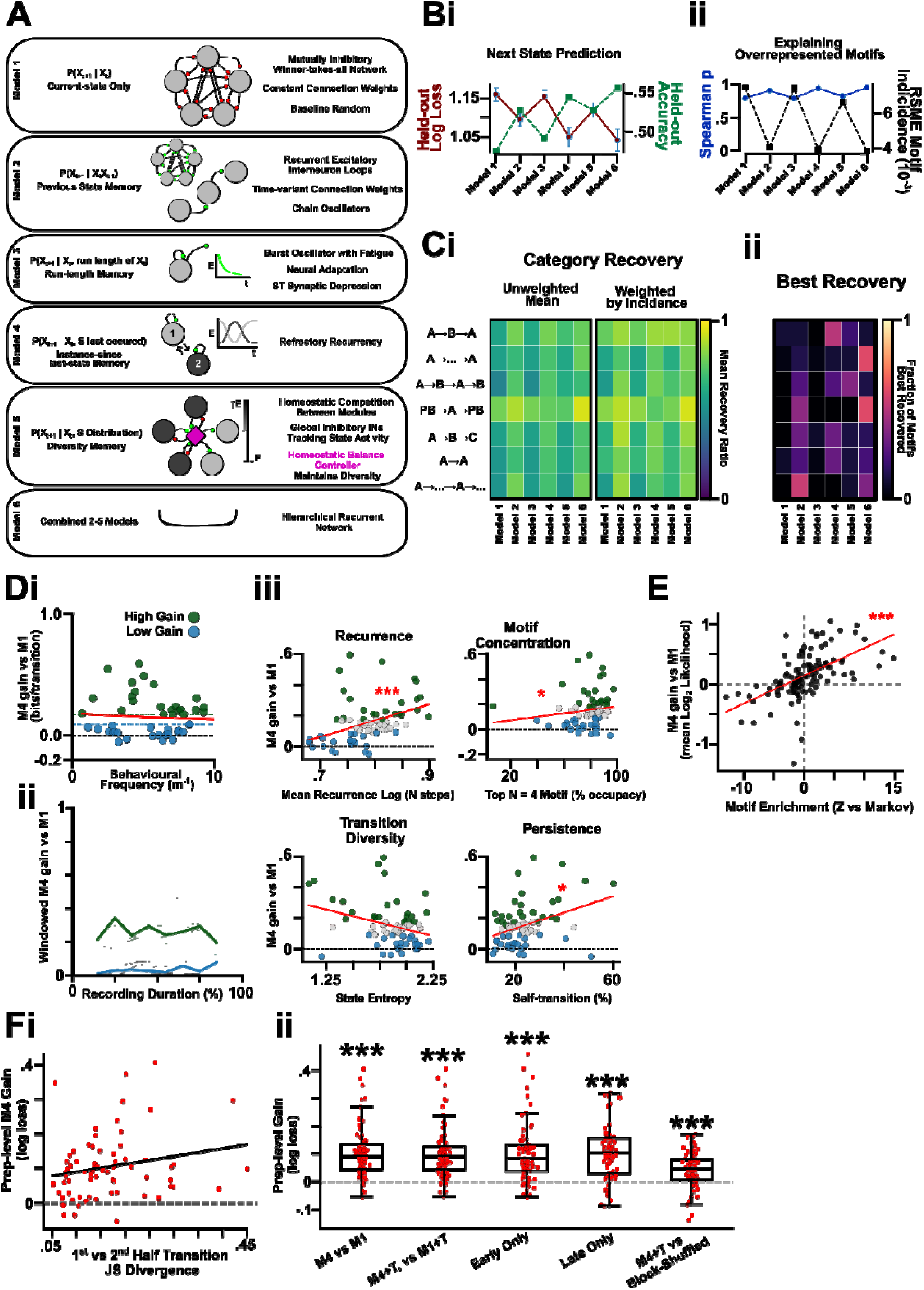
A Since-last-state Model Best Predicts Fictive Sequence Evolution and Captures Overrepresented Motifs. (**A**) Probabilistic models featuring a variety of prior state dynamics were considered to explain multi-step fictive evolution captured by a variable-order Markov chain. (**B**) Quantified predictivity of the probabilistic models for (**i**) next state prediction and (**ii**) capacity of each probabilistic model to explain the overrepresented motif. (**Ci**) Recovery of each specific motif categories by each probabilistic model and (**ii**) which model best recovers and predicts motif categories. The motif categories shown include: three-step motif where the a type of fictive event starts the motif and that state reappears after an intermittent fictive event (A→B→A), fictive motif where the 1 fictive event is the same as the final fictive event (A→…→A), motifs that alternative between two different fictive events (A→B→A→B), motifs that start and end with a posterior burst (PB→A→PB), a chain of different fictive events (A→B→C), motifs which have adjacent repeating fictive elements (A→A), and motifs with mixed repeating chains (A→…→A→…). (**D**) Since-last-state memory is (**i**) invariant to the frequency of events in the recording period and exists most strongly a subset of preparations (N = 29, defined by bits/transition threshold window of > 0.2) where model 4 offers a high explanatory gain with (**Dii**) since-last-state memory present across the recording period in those high-gain preparations. (**Diii**) Model 4 uniquely captures Recurrence, motif concentration, transition diversity, and persistence of certain overrepresented fictive motifs which drives its explanatory power over the baseline model 1. (**E**) The most frequent multi-step fictive motifs that are enriched in the variable-order compared to primary Markov model are proportionally captured by model 4 relative to the baseline model 1. (**Fi**) Preparation-level M4 gain, quantified as the reduction in held-out log loss relative to M1, plotted against first-versus second-half transition JS divergence. Each point represents one preparation. Linear fit is shown; asterisk indicates a significant Spearman correlation. (**ii**) Preparation-level memory gain across drift-control analyses. M4 improves prediction over M1 in the full dataset, after temporal-position control, within early and late recording halves, and relative to a block-shuffled since-last-state control. Red points indicate individual preparations; boxplots show median and interquartile range

To determine whether the weaker performance of recent-history models depended on the chosen recent-window size, we repeated model 5 and model 6 fitting across recent windows ranging from 2 to 30 events. Across all window sizes, model 5, which incorporates recent-window balance and diversity, produced only small gains over the current-state model, with median gains ranging from −0.001 to 0.009 bits/transition. Such remained far below the baseline since-last-state gain of 0.131 bits/transition. Thus, recent-window diversity or balance alone did not explain the latent fictive memory hysteresis. The full-memory model remained the best-performing model across window sizes, but only marginally exceeded model 4, with median M6-over-M4 gains ranging from 0.008 to 0.016 bits/transition. Consequently, model 4 captured the dominant memory signal across all recent-window choices, indicating that the principal predictive feature reflects state-specific recency rather than arbitrary recent-window composition. Model 4’s predictive advantage was also robust across logistic-regression regularisation strengths and alternate since-last-state encodings. Across raw, log-transformed, and capped recency predictors, the since-last-state model retained a positive preparation-level gain over the current-state model (Figure 4C), with representative median gains of 0.13–0.15 bits/transition and significant paired improvements across preparations.

Since-last-state memory appeared largely independent of overall activity rate, instead being driven by categorical features of fictive sequence organisation. The relative explanatory gain of model 4 was not related to the frequency of fictive activity across preparations (Figure 4Di). Consistently, high-gain and low-gain preparations did not differ in event frequency (high-gain median = 6.27 events/min; low-gain median = 6.13 events/min) or total sequence length (high-gain median = 282 events; low-gain median = 276 events). Instead, preparations separated into distinct predictive regimes, with high-gain preparations showing substantially greater benefit from since-last-state memory than low-gain preparations (median M4 gain: high-gain = 0.236 bits/transition; low-gain = 0.029 bits/transition; q < .001). In high-gain preparations, the explanatory benefit of since-last-state memory persisted throughout the recording rather than appearing as a transient or isolated epoch (Figure 4Dii). Across windowed preparations, M4 gain remained positive across the recording period, with average windowed gains ranging from 0.114 to 0.185 bits/transition across time bins, and preparation-level gain slopes were not significantly different from zero. Thus, since-last-state memory is not simply a brief phase-specific phenomenon, but a persistent feature of fictive dynamics in preparations where this form of history dependence is expressed.

The explanatory power of model 4 was associated most strongly with recurrence-based sequence features. Preparations with greater M4 gain showed higher probability of returning to a recently expressed state within four events (ρ = 0.485, q = .004), within three events (ρ = 0.403, q = .005), and within two events (ρ = 0.320, q = .030). Conversely, M4 gain was negatively associated with mean recurrence lag (ρ = −0.350, q = 0.018), indicating that high-gain preparations tend to revisit states more quickly. There was also a significant association with persistence, with M4 gain positively correlated with self-transition fraction (ρ = 0.305, q = 0.0265) and negatively correlated with switching fraction (ρ = −0.305, q = 0.0265). By contrast, broader diversity and motif-concentration metrics did not survive FDR correction, including transition entropy, state entropy, top trigram fraction, and repeated quadgram fraction. Therefore, high-gain since-last-state memory is not best described as a simple loss of behavioural diversity or as the emergence of one specific motif class. Rather, it reflects a constraint on categorical recurrence: recently expressed fictive states remain more available for re-entry over the next few fictive behavioural events.

Importantly, model 4 captured not only the presence of recurrent motifs but also their degree of enrichment relative to a first-order Markov null (Figure 4E). Across Markov-deviant motifs, motif-level M4 gain was strongly correlated with motif enrichment magnitude (main motif set: ρ = 0.657, p < .001 strict motif set: ρ = 0.761, p < .001). Thus, motifs that deviated more strongly from first-order Markov expectations tended to be those whose prediction was most improved by since-last-state information. Such indicates that non-Markovian sequence structure arises from a history-dependent process rather than stochastic or memoryless dynamics. Consistent with this, model-based motif recovery showed that the full memory model best recovered the largest number of deviant motifs (70 motifs), but the since-last-state model was the next strongest model (49 motifs), exceeding the previous-state, recent-diversity, and run-length models. Thus, state-specific recency captures a major component of the motif structure identified in the previous analysis.

Given self-recurring motifs, such as A→B→A, were overrepresented in fictive dynamics, we next confirmed that since-last-state memory promotes recent-state Recurrence rather than recent-state suppression or refractory dynamics. For each candidate state, we calculated the probability that this state would occur next as a function of the number of intervening events since that state was last expressed. Across states, recently expressed fictive behaviours were more likely to recur. In the raw analysis, recent-minus-distant recurrence was positive for all fictive states: AB +0.096, AS +0.144, BW +0.129, FW +0.039, and PB +0.140, with all FDR-corrected q-values significant. Importantly, this recent-state enrichment remained after excluding immediate self-repeats, after excluding rows in which the candidate state was already the current state, and under a conservative analysis excluding both forms of local persistence. In this conservative analysis, recent-minus-distant recurrence remained positive for all states: AB +0.133, AS +0.229, BW +0.302, FW +0.061, and PB +0.133, with all q-values ≤ .001. Thus, the since-last-state signal is not reducible to adjacent self-transition or current-state persistence. Instead, it reflects a broader categorical recurrence memory in which recently expressed states remain transiently available for re-entry.

Conceptually, the since-last-state effect is consistent with an adaptive winner-takes-all organisation of competing motor programs. In a classical winner-takes-all circuit, the currently dominant motor module suppresses alternative modules to stabilise behavioural output [22], [23], [24], [34]. However, the recurrence-promoting memory observed here suggests that ‘winning’ fictive states are not simply reset or made refractory after expression. Instead, recently expressed motor programs remain transiently more available for re-entry, effectively biasing future competition without fully locking the system into repetition. Thus, fictive sequence evolution may reflect *adaptive* winner-takes-all dynamics in which recent winners leave a categorical trace that reshapes the competitive landscape, promoting structured recurrence and alternation between motor programs while preserving access to alternative fictive outputs.

Given non-stationarity can generate apparent higher-order sequence structure, we next evaluated whether since-last-state memory remained predictive after accounting for temporal drift in fictive sequence dynamics (Figure 4F). Despite adding temporal predictors describing each transition’s relative position in the recording, since-last-state memory retained improved prediction over time-controlled current-state models. Model 4 improved prediction over the current-state model at the preparation level (median gain = 0.0906 log-loss units, p < .001) and this effect remained after adding time quartile predictors to both models (median gain = 0.0908, p < .001) or time decile predictors (median gain = 0.0920, p < .001). Since-last-state memory also improved prediction within both early and late recording phases, with median gains of 0.0825 log-loss units in the early phase (p < .001) and 0.1026 in the late phase (p < .001). Finally, we performed a block-shuffled control in which since-last-state predictors were shuffled within each preparation and temporal quartile. Such preserved preparation identity, recording-phase structure, and the local distribution of since-last-state values while disrupting their event-wise alignment with the next transition. The real time-controlled since-last-state model outperformed this block-shuffled control (median gain = 0.0439 log-loss units, p < .001) demonstrating that the predictive signal depends on precise categorical sequence history rather than non-stationary drift alone.

The magnitude of since-last-state gain was weakly but significantly associated with first/second-half transition drift, both before temporal control (ρ = 0.240, p = 0.047) and after temporal control (ρ = 0.245, p = 0.047). Thus, preparations with greater slow reconfiguration tended to show stronger local event-history dependence. The recency signal was not eliminated by temporal controls, indicating that it is not simply an artefact of slow non-stationary drift in fictive transition or frequency properties. However, its association with transition drift suggests that local categorical memory and gradual reconfiguration of the fictive landscape may be coupled rather than independent properties of spontaneous motor network dynamics. Overall, categorical since-last-state recency explains fictive transition structure beyond the non-stationary drift exhibited by the fictive network.

Taken together, fictive behavioural transitions are strongly shaped by adaptive dynamics that depend on how recently a state was last expressed within the sequence. Since-last-state memory persists after temporal controls, but its magnitude covaries with drift, suggesting that local history dependence and slow landscape reconfiguration may be coupled features of the same evolving fictive network. Importantly, since the sequence information is categorical, devoid of relative activity timing, and fictive rhythms have varying periodicity [1], [31], [33], this suggests an event-based rather than time-based hysteresis.

### Latent Network States Organise Fictive Activity Around AS-Dominated Transition Regimes

Observable fictive behaviours are structured by distal transition hubs, multi-step motif memory, and slow non-stationary drift. However, each detected fictive behaviour is only an observable readout of an underlying network state; the same fictive behaviour may arise from different latent network configurations with different persistence and future transition probabilities. We therefore asked whether spontaneous fictive behavioural sequences could be decomposed into latent fictive network regimes using hidden Markov (HMM) and hidden semi-Markov (HSMM) modelling.

HMM model selection supported a six-state latent representation of fictive sequence structure (Figure 5A). However, dwell-time analysis showed that several latent regimes were not well described by memoryless geometric persistence, indicating that the probability of leaving a latent state was not constant over time (Figure 5B,C). We therefore implemented an observable HSMM in which state persistence was modelled explicitly. Such produced a six-macrostate, nine-substate representation of fictive dynamics, allowing latent regimes to differ both in the behaviours they emitted and in how persistently they held the network within a particular configuration (Figure 5D). Thus, latent fictive states are not simply instantaneous labels for observed behaviours, but persistent network regimes with distinct dwell and transition structure.

**Figure 5.**
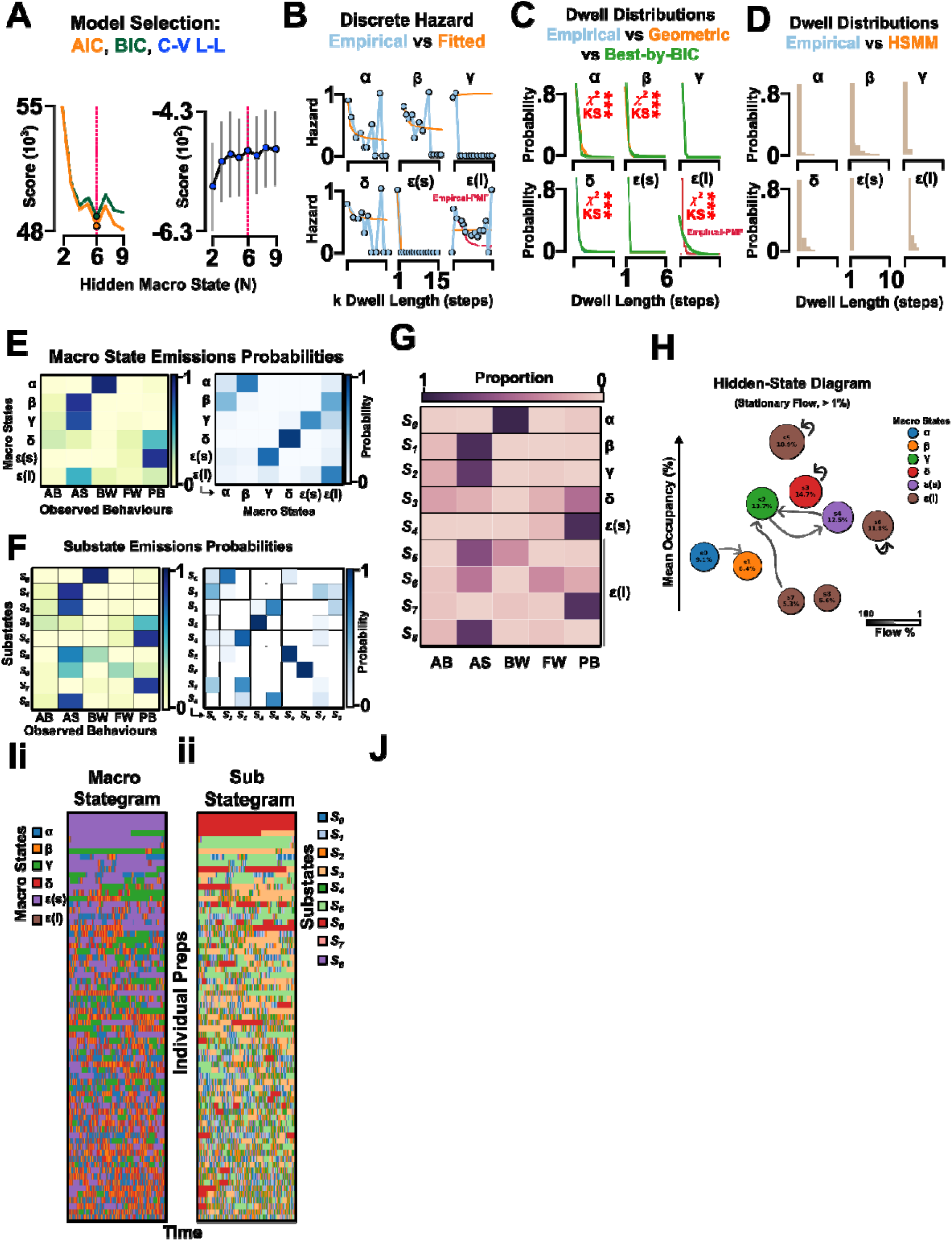
Hidden Semi-Markov Model Reinforces Anterior Asymmetric as Central Hubs in Wave vs non-wave Dynamics. (**A**) Model selection to determine the number of hidden states through AIC and BIC score and cross-validated log-likelihood score. Red dashed line indicates AIC, BIC minimum and log-likelihood maxima indicating 6 hidden states (α, β, γ, δ, l2(s), l2(l)) within the fictive spontaneous data. (**B**) Discrete hazard (i.e., the instantaneous probability that the network transitions out of its current hidden state) for dwell length for empirical and fitted model data segregated by hidden state. The bifurcated lll hidden state was non-geometrically distribute thus PMF hazard was fitted (red line). (**C**) Dwell distribution comparisons between empirical (blue), geometric (orange), and best-by-Bayesian information criterion (BIC) score (green) for each hidden state. The standard HMM assumes geometrically distributed dwell durations because the probability of exiting a hidden state is constant at each event. State-specific dwell analysis showed that one AS-associated state, ε, violated this assumption by containing both short-dwell and long-dwell components. Thus, ε is represented as two HSMM substates, ε(s) and ε(l), separated by the empirical dwell-duration cutoff used in the Methods. In the HSMM, state persistence is generated by explicit dwell-duration distributions rather than by HMM self-transition probabilities. (**D**) The probability of each hidden macro state’s dwell length for hidden semi-Markov chain (HSMM) simulation (orange) compared against empirical data (blue). (**E**) Heatmaps for the emission probabilities of each macro state and the observed fictive behaviours represented in the macro state (left) and transition between macro states (right). (**F**) Heatmaps for the emission probabilities for each substate and the observed fictive behaviours represented in the substate (left) and transitions between substates (right). (**G**) The proportion of observed fictive behaviours being represented in each macro and macro-associated substate. (**H**) The hidden-state stationary flow diagram between the substates with their macro states labelled by node colour. (**I**) The time normalised (%) occupancy of each individual preparations (N = 69) in their (**i**) macro states and (**ii**) hidden substates. (**Ji**) Infographic representing the nuanced a wave (W) – distal (NW) activity transition dynamic axis showing which macro and substates are conceptually within this axis and the dominant (> 20%) transitions between each substate shown. Green arrows indicate self-reinforcing states for either W or NW behaviours. Purple arrows indicate transitions from W → NW states (dashed) or NW → W (whole). (**Jii**) The mean and ranged normalised occupancy % of each substate across all preparations. (**Jiii**) The normalised % difference in transitions from W minus to NW-related substates with error bar swarm indicating population-level equivalency in W↔NW transitions despite some individual-level bias.

The HSMM emission structure revealed biologically interpretable latent regimes rather than arbitrary mixtures of fictive behaviours (Figure 5E,F). Several states were dominated by distal activity, particularly AS and PB, whereas others combined distal activity with wave-related outputs such as BW or FW. Such organisation mirrors the observable transition analysis, where AS and PB formed a central distal transition subspace. Importantly, AS was emitted across multiple latent regimes, indicating that AS fictive activity is not a single homogeneous behavioural state. Instead, the same scored AS event can occur as part of an AS/BW wave-access regime, an AS/PB distal regime, or an AS→AB/PB transition regime. Thus, the HSMM resolves the observable behavioural landscape into hidden configurations with distinct emission, persistence, and transition properties.

Latent transitions were also structured rather than random. Hidden-state transition matrices and flow diagrams showed preferred routes through latent-state space, with AS-rich and AS/PB-rich regimes acting as major conduits between distal and wave-related dynamics (Figure 5E–H). Such organisation was not dependent on a single visualisation or edge-weighting choice: probability flow, symmetric flow, surprise flow, mutual-information flow, and macrostate flow all recovered broadly comparable hidden-state structures, with states positioned by their mean occupancy across preparations (Supplementary Figure 3). Thus, the inferred latent map reflects a robust transition organisation in the data, rather than an artefact of one graph metric.

The substate decomposition further showed that similar observable behaviours can arise from latent contexts with different local sequence rules. Across macrostates, state entropy, self-transition fraction, switch fraction, recurrence lag, and transition entropy all differed strongly (state entropy: χ² = 91.57, p < .001 self-transition fraction: χ² = 91.04, p < .001; mean recurrence lag: χ² = 90.17, p < .001; transition entropy: χ² = 77.26, p < .001). Similar differences were present across hidden substates, including mean recurrence lag, return-within-2, and state entropy (mean recurrence lag: χ² = 35.90, p < .001; return-within-2: χ² = 31.89, p < .001; state entropy: χ² = 31.77, p = .001). Therefore, scoring the same observable behaviour does not necessarily identify the same latent network condition. An AS event, for example, can reflect a stable distal regime, a recurrent AS/PB loop, or a transitional context linking distal and wave-related activity.

Posterior predictive and robustness checks showed that the HSMM captured several major empirical features of fictive sequence organisation while also identifying its limits (Supplementary Figure 4). The model reproduced broad occupancy, dwell, entropy, and event-level transition structure. Dwell fits were strongest for BW, AB, and PB, whereas AS dwell structure was the most difficult feature to reproduce. Explicit AS splitting improved this mismatch, with the AS-split model outperforming a negative-binomial AS dwell model for AS dwell reconstruction (AS Wasserstein: AS-split = 0.108; AS-NB = 0.215), and a q = 0.75 split giving the lowest average dwell mismatch across the tested robustness grid. Under this split, permutation KS tests did not reject empirical-versus-simulated dwell distributions for any observable state after FDR correction (all KS q ≥ 0.456), although residual chi-square differences remained for FW and AS. Thus, explicit dwell modelling and AS substate splitting substantially improved the HSMM fit, but AS-rich regimes retained residual complexity not fully captured by a time-homogeneous latent-state model.

However, the HSMM did not fully explain the higher-order motif structure identified earlier. To directly compare the HSMM with the other sequence models, we evaluated how well model-generated sequences reproduced the empirical motif-enrichment landscape (Figure 6). Such separated two related questions: whether a model reproduced motif occurrence, and whether it reproduced motif deviancy relative to a first-order Markov null. The variable-order Markov model provided an upper-bound reconstruction of local sequence dependence, reproducing empirical motif-enrichment scores almost perfectly for both tri-grams and quad-grams (3-grams: ρ = 0.999, direction agreement = 98.4%; 4-grams: ρ = 0.998, direction agreement = 97.6%) (Figure 6A,B). Among interpretable models, the since-last-state model gave the strongest account of motif organisation, outperforming both the first-order Markov model and HSMM in reproducing Markov-deviant motif enrichment (3-grams: ρ = 0.796, direction agreement = 84.4%; 4-grams: ρ = 0.768, direction agreement = 85.1%) (Figure 6A,B). By contrast, the HSMM captured broad motif occurrence but only moderately reproduced motif enrichment. HSMM-simulated motif counts were strongly correlated with empirical motif counts for both 3-grams and 4-grams (3-grams: ρ = 0.943; 4-grams: ρ = 0.849), indicating that the model captured much of the occupancy and transition structure of the sequence. However, HSMM motif-enrichment scores showed only moderate correspondence with empirical enrichment scores (3-grams: ρ = 0.419, direction agreement = 57.4%; 4-grams: ρ = 0.457, direction agreement = 62.8%) (Figure 6D). Such distinction was strongest for recurrence-like motifs, where the since-last-state model reproduced recurrent motif structure substantially better than the HSMM, particularly for 4-grams (M4: ρ = 0.744, direction agreement = 89.0%; HSMM: ρ = 0.409, direction agreement = 66.4%) (Figure 6E). Thus, the HSMM captures which latent regimes the system occupies and how they persist, whereas since-last-state memory better captures which multi-step routes are revisited, amplified, or suppressed.

**Figure 6.**
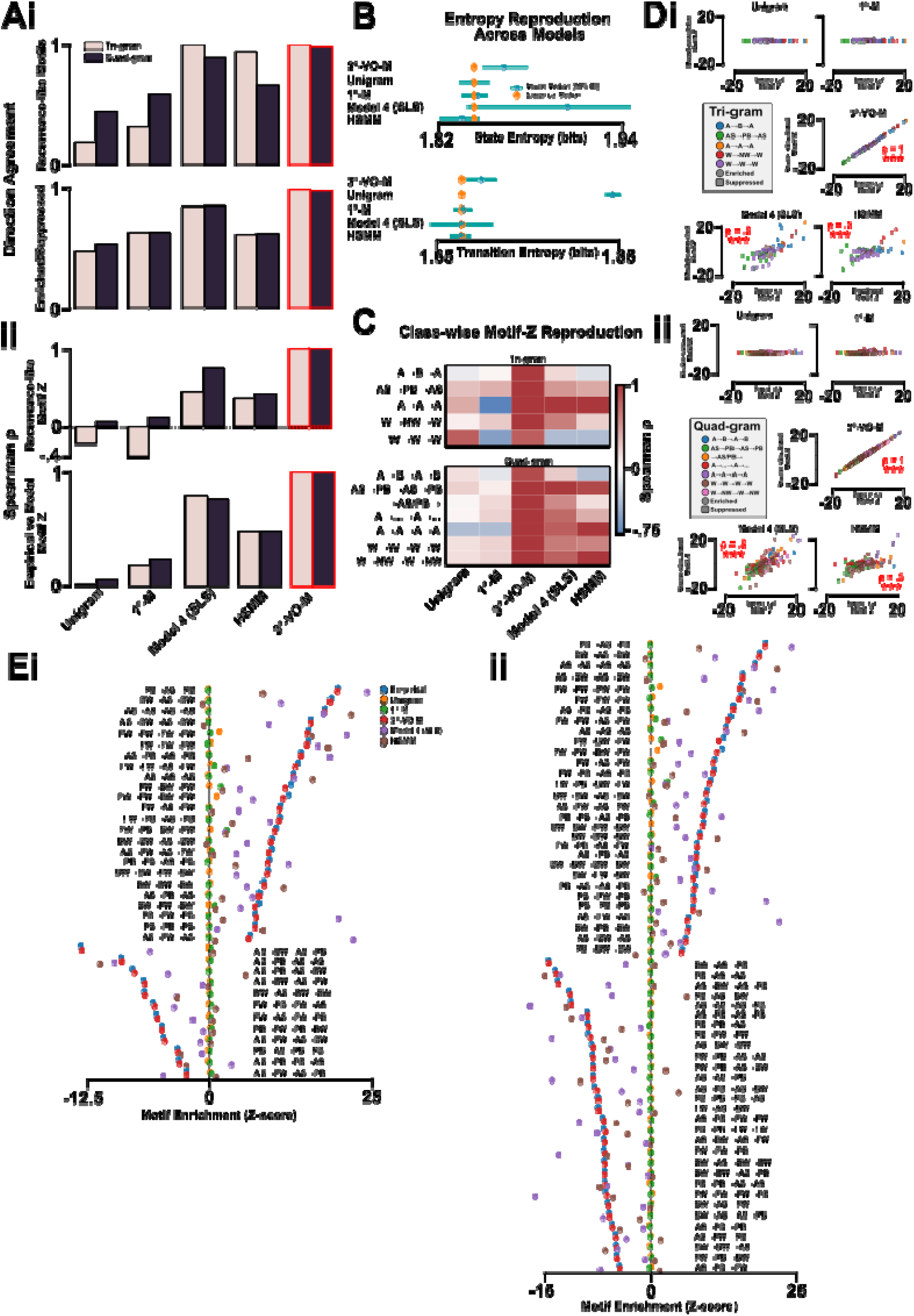
Motif-structure Comparisons between HSMM, Unigram, first-order Markov, Variable-order Markov, and Since-last-state Memory Models. (**Ai**) Direction agreement and (**ii**) spearman correlation of recurrence-like motifs, status (enrichment/suppressed) relative to empirical data, z-score of recurrence-like motifs and empirical vs model for unigram, primary Markov model (1°-M), since-last-state model 4 (SLS), hidden semi-Markov model (HSMM), and 3-step-variable-order-Markov model (3°-VO-M) of multi-step trigrams and quadgrams. The 3°-VO-M represents a perfectly-fit model for empirical data as the model is directly trained to generate empirical trigrams and quadgrams. (**B**) The entropy reproduction across all models for state and transition dynamics. (**C**) Spearman correlation of class-wise motif z-score reproduction across the models. (**D**) Relative motif z-score compared against empirical z-scores of (**i**) trigrams and (**ii**) quadgrams for each model. Spearman correlation and statistically significant relationships reported. **(E)** Relative enrichment or suppression of (**i**) recurrence-like motifs and (**ii**) model reproduction for each model. Note that since-last-state model 4 captures empirical relative enrichment and suppression stronger than HSMM, but both consistently stronger than unigram or 1°-M model.

Together, the HSMM and since-last-state analyses provide complementary explanations of fictive rhythmogenesis. The HSMM identifies the latent regimes through which the fictive network moves, demonstrating that observable behaviours—especially AS—can be generated from multiple hidden contexts with distinct transition and dwell properties. In contrast, the since-last-state model identifies the event-history rule that biases movement through this latent landscape, explaining why particular recurrent motifs are enriched beyond first-order Markov expectations. Thus, fictive motor dynamics are best understood as the interaction between latent regime structure, AS/PB-centred hidden-state flow, slow non-stationary drift, and categorical recent-state memory. The final model reflects how the isolated fictive network can maintain structured motor diversity without collapsing into either random exploration or rigid repetition.

## Discussion

In the larval locomotor system, spontaneous fictive activity is structured by interacting mechanisms operating at multiple levels of sequence organisation. We show that fictive locomotor transitions are shaped by since-last-state memory which biases the network towards repeating recently executed sequences of motor programs. The hysteresis in fictive transitions is not explained by first-order transitions, run length, recent repertoire diversity, self-persistence, nor non-stationary drift. Instead, recently-expressed states remain transiently available for recurrence, producing structured higher-order motifs within a slowly drifting fictive landscape. However, 1^st^-order and higher-order Markov analysis reveals that fictive locomotor activity also occupies distinct coordination regimes, which bias transition probabilities towards more diverse output states, without necessarily altering the observable motor programs. Within this organisation, anterior asymmetric activity – corresponding to larval head sweeps – acts as a transition hub linking multiple locomotor states and promoting output diversity. Taken together, the *Drosophila* fictive network appears to recall and promote diversity through specific, consistent, and identifiable motif structures determinable through hidden-state analysis.

### Fictive Dynamics Maintain Flexibility without Instability

The emergence of diverse fictive motor programs with constrained, non-random higher-order structure suggests that rhythm-generating networks in *Drosophila* larvae do not simply explore motor space randomly. Instead, interacting CPGs appear to contain latent transition rules that organise the transition between motor programs. Canonical work within rhythm generation across invertebrate and vertebrates demonstrated how multiple rhythmic outputs evolve from shared or interacting circuits that are persistently shaped by neuromodulation, intrinsic state, and circuit history [4], [7], [8], [10], [11], [12], [13], [35], [36], [37], [38]. Here, we extend this view by identifying specific organisational features within isolated *Drosophila* fictive locomotion that suggest fictive networks evolve through structured regions of motor space in the absence of sensory feedback or reafferent correction. A core problem of multifunctional motor systems is the need to generate multiple antagonistic outputs whilst avoiding becoming trapped in one output regime: maintaining flexibility without instability [5], [17], [35], [36]. The intrinsic features we identify act as mechanistic routes back to a diverse, transition-ready state after periods of induced, or maladaptive bias. Specifically, exploratory behaviours, like *Drosophila* anterior asymmetric activity, may operate as an intrinsically-primed hub that links otherwise distinct motor programs and helps restore the network to a diverse, transition-ready state after biased activity. Conceptually, this organisation would speak to the functional hierarchy between different CPGs where those circuits responsible for exploratory behaviours are potentiated after bias and consequently potentiate alternate CPG in the absence of sensory reafference. Indeed, this interpretation is consistent with our previous work which demonstrated anterior asymmetric activity was proportionally promoted after sustained octopaminergic-induced fictive forward bias [3]. Importantly, the HSMM states reported here are not intended as one-to-one behavioural labels, but as latent contexts that bias the probability of subsequent fictive events, analogous to internal-state models in computational ethology [39].

### Putative Neural Architectures for Hubs & Time-invariant Memory

The structure transition patterns and constraints on diversity we report here produces constrained testable hypotheses for CPG motifs architecture. Fundamentally, the results here raise an interesting central question: how can a time-dependent biological process generate event-count memory? Here, the model demonstrates that memory is most visible during motor program coordination, but not that motor program output itself is the biophysical coordinate of memory. The central constraint is that the fictive system is oriented towards structured diversity, not randomness – at the global level, sequence transitions are organised to hubs of high entropy, but at the local sequence level, since-last-state memory constrains behavioural dynamics to promote short-term recurrence.

The circuitry underlying motor output must maintain a time-invariant, short-term trace of recent activity to adapt transitions based on how recently fictive behavior has recently occurred. Specifically, recently expressed fictive states were more likely to reoccur, demonstrating the fictive system retain a short-term categorical trace that transiently increases the availability of recently-occupied states. Conceptually, residual network excitability, excitatory recurrent circuit motifs, and activity-dependent adaptation mechanisms like spike-frequency adaptation, Na^+^/K^+^ pump currents or short-term synaptic facilitation [19], [40], [41], [42], [43] may underpin assuring output stabilisation while not impeding potentiating output diversity. Importantly, the manifestation of these mechanisms will diverge depending on if CPGs are independent or interlocked for distinct fictive outputs [7]. In effect, the activation threshold for recently-active motor circuits for one fictive behaviour may become temporarily altered reminiscent of locomotor adaptation currents in CPG networks [44], [45]. Alternatively, short-term memory may be encoded by an “integrator” neuron that accumulates activity from a motor module and alters reactivation [46], [47], [48]. We would expect the decay in the integrator neuron’s activity to correlate to the timescale for the memory. Similar circuit motifs integrating activity-dependent recurrence are found in tadpole locomotor networks [49], [50], circuits in the central complex of *Drosophila* with state-linked activity patterns [51], [52] state integrators in STG circuits [6], [7], and mammalian spinal circuits where interneurons gate locomotor states subject to modulatory processes [12]. In effect, the global organisation of *Drosophila* fictive rhythms as described here suggest that such integrator and activity-adaptive circuits are fundamental and essential to promotion of motor program diversity.

Our hub-like dynamics that precede diverse outputs may speak to a common interneuron population between components of a CPG responsible for executing distinct fictive outputs. Such a “hub” neuron/circuit that selectively excites or inhibits modules of the CPG is common in many insect motor systems in the form of descending command neuron circuits [53], [54], [55]. Indeed, interneurons which coordinate activity between segmental oscillators in invertebrates [8], [56], [57], could form the basis of such a “hub” neuron due to their high connectivity across modules.

Previously identified components of the larval locomotor circuit may have activity that tracks the general latent state of the network that persist beyond individual fictive behavioural output. We would predict that global activity signatures across different interneuron subsets linked to CPGs might represent internal network configurations. Consequently, wide-scale adaptations to CPG global activity signatures could be used to bias the latent state of the network, creating predictable alternations in transition patterns. For example, intrinsic alteration of *Drosophila* PMSI interneurons activity would bias wave propagation dynamics [58], modulatory input to A27h neurons could bias the system towards a wave-like regime [59] whereas GDL interneuron modulation could bias the network to distal activity regimes [60] and command neurons (e.g., MDN) could directly shift the network into a wave regime [53]. These specific neurons – or generally similar motifs – could be the site of monoaminergic modulation responsible for locomotory gating (similar to MCN1 reconfiguration of STG circuits) [3], [17], [61], [62], [63]. Indeed, we can conceptualise hidden states as an ensemble of cellular network, and network properties that alter CPG circuits to generate similar fictive outputs but with different transition biases. Indeed, neuromodulation may play a critical role in functionally modifying components in the motor system to shift the internal hidden state. This could conceivably arise from modulation of neuronal intrinsic properties and synaptic dynamics which have been shown to dramatically alter the fictive landscape of motor systems (e.g., octopamine/tyramine [3], [64]; dopamine [65], [66]) [17], [67].

The gradual shifting of the fictive dynamic structure over time points to slowly-changing variables within the motor circuit. Specifically, we would expect periodic ionic homeostasis sufficient to alter network excitability in the core components of the circuit motif involved in action selection, motor competition, and the CPG network at large. Indeed, intrinsic monoamine signalling [17] and activity-dependent metabolism [12], [15], [50] alter circuit performance and network dynamics putatively responsible for the gradual shift in fictive structure. Conceptually, shifts in rhythmogenic activity may represent a feature to ensure adaptive behaviour. For instance, gradual changes in extracellular potassium can alter state transitions due to bistable firing regimes [68]) with a underexplored role in ion homeostasis due to glia being a novel avenue for exploration [69].

A curious feature exhibited by our since-last-state memory analysis is that the intrinsic memory appears invariant to the frequency of activity within a preparation recording and, more importantly, is revealed purely through sequence information, not the relative incidence timings of those fictive behaviours. Thus, any putative architecture encapsulating since-last-state memory must be time-invariant and sensitive uniquely to the categorical instances of fictive activity, not their relative timing. This memory mechanism existed throughout all latent network states with variability only emerging on an inter-preparation, not inter-hidden state, basis - implying a global adaptation regime applied across different circuit configurations. Consequently, any of the aforementioned mechanisms must include a global, time-invariant, and state-invariant mechanism where the neural network responsible for tracking and selective enforcing diversity must hold an evolving impression of the fictive diversity without meaningful decay and thus loss of information over time. A clear parallel for since-last-memory can be drawn to work deciphering the neural correlates of time-invariant mental representations within working memory in the mammalian neocortex [70], [71]. Interestingly and informatively, modelling efforts to encapsulate working memory dynamics at a neuronal level demonstrate the canonical time-invariant features of neural activity are not incompatible with fixed, time-invariant representations. Indeed, by using recurrent connectivity with time decay under the Feature Vector REcombination (FEVER) principle, a neural network can preserve encoded features (e.g., mental representations, sensory information) over time [72] showing the biological plausibility of such a putative core feature in *Drosophila* fictive dynamics.

Experimentally, we propose a few ways to test the results reported here. Firstly, if anterior asymmetric rhythms are a true transition hub, perturbing asymmetry-associated circuits (e.g., thoracic GABAergic neurons [1]) should reduce outgoing transition entropy and reduce access to alternative fictive trajectories. Secondly, if since-last-state memory reflects recurrent priming, perturbing recurrent excitation or short-term synaptic facilitation should weaken ABA/ABAB-specific motif enrichment from the fictive system. Thirdly, if since-last-state memory is a global state variable – as interpreted through the HSMM – then the inter-preparation gain in since-last-state memory explanatory power should covary with global or specific network activity (i.e., calcium levels).

### Translating Fictive Dynamics to Intact Behaviour

Fictive dynamics are closely reminiscent of intact behaviour [31], [33] implying the hallmarks of hierarchical transition dynamics reported here should be discoverable in intact, freely-moving *Drosophila* larva. We would predict that headsweep behaviours to remain the highest entropic state and there to be robust behavioural trajectory between behaviours related to exploration (i.e., headsweeps) versus behaviours related to sustained movement (i.e., crawling). Indeed, within freely-moving adult *Drosophila* and mouse behaviour a complex array of distinct movements can be decomposed into a distinguishable set of stereotyped behaviours [73], [74], [75], [76] implying our analysis here could be generally informative to a variety of animal locomotion. Intuitively, locomotor CPG networks would be biased to fundamental exploratory behaviours; however, rhythm-generating circuits would also potentiate all possible post-exploratory behaviours, as that could maximise behavioural adaptability. Specifically, *Drosophila* headsweep behaviour is primarily a scouting behaviour for a variety of external stimuli: predation, food, light, navigation, avoidance of aversive stimuli [2], [77], [78], [79], [80], [81]. Thus, we would expect CPG networks to be organised to potentiate exploratory-like behaviours, then decrease the barriers of entry into all other fictive patterns.

While fictive dynamics exhibit the same categorical behavioural types as intact behaviour, freely-moving *Drosophila* larva show a biased towards forward and headsweep locomotion whereas fictive locomotion exhibits a higher incidence of fictive backward activity [31]. To our knowledge, the type of analysis above has yet to be applied to intact *Drosophila* behaviours. We predict that the fundamental system of exploratory-bias followed by post-exploration diversity to hold true at the behavioural level. Across many model systems, there is considerable evidence of sensory-gating of motor state transitions [79], [80], [81], [82], [83], [84]. Even if the biased in the state transition hierarchy is distinct between intact and fictive preparations, we nonetheless would expect the same underlying mechanistic rules to be conserved. Regardless, any divergency between fictive and intact dynamics is informative in understanding sensory integration by providing a reference on how the central rhythmogenetic system is modulated by afferent sensory information.

### Adaptive Winner-Takes-All Competition Amongst Central Pattern Generating Networks

The observed competition between distinct larval CPG components points towards a mechanism combining mutual inhibition with a biological constraint of adaptation or fatigue. Conceptually, competition between neural networks can be encapsulated as a winner-takes-all circuit whereby a system with the highest excitation, and thus greatest capacity to inhibit competing circuits, dominate and execute an output [85], [86], [87]. Here, the existence of instance-since-last-state memory suggests that recently active motor modules become transiently supported enabling a given modules to retain the opportunity to dominate output. Thus, rather than a canonical winner-takes-all motif, we conceptualise inter-CPG competition in the *Drosophila* larva as *adaptive* winner-takes-all competition.

An *adaptive* winner-takes-all model of fictive rhythmogenesis therefore makes several testable predictions. Perturbing mechanisms that encode recent motor history, such as activity-dependent ionic currents, recurrent excitation, or inhibitory motifs, should directly alter the probability of re-entering recently active fictive states. In addition, if competing CPG modules are organised into stable coordination regimes, weakening mutual inhibition should reduce regime separation and increase mixed or co-expressed fictive outputs. Likewise, the dwell duration of wave-like or lateralised regimes should remain stochastic but structured, with blocking adaptation predicted to prolong dominance and flatten state-exit hazard rates. Transitions between competing regimes should also pass through specific intermediate or hub states, such as anterior asymmetric activity in *Drosophila*; silencing neurons that support this hub motifs should impair switching between otherwise exclusive motor programs. In parallel, slowly varying internal variables, including neuromodulatory gain or synaptic weight changes, should bias hidden regime occupancy without necessarily changing the immediately observable fictive pattern. Finally, when the network occupies high-entropy boundary states between competing attractors, brief sensory or optogenetic perturbations should be especially effective at redirecting the system toward whichever motor module is most excitable or least adapted. Together, these predictions distinguish *adaptive* winner-takes-all competition from a purely stationary latent-state model by requiring experimentally manipulable links between inhibition, adaptation, hidden-state occupancy, and history-dependent transition bias. Future experiments that induce direct perturbation of candidate inhibitory, or adaptation mechanisms will help further refine the mechanistic underpinnings of competitive interactions in our system and others like it.

Overall, we show the *Drosophila* fictive network exhibits two mechanisms to regulate output diversity. The network promotes output diversity by promoting anterior asymmetric activity that evolves into a more diverse repertoire of fictive activity. Simultaneously and antagonistically, the network can restrain output diversity through promoting recurrency of recently-explored states. These competing mechanisms are novel features of *Drosophila* fictive rhythmogenesis important for understanding how motor systems regulate output diversity.

## Methods

### Animal Rearing, Dissection, and Calcium Imaging

Imaged *Drosophila melanogaster* 3^rd^ instar larvae were reared in vials containing cornmeal-based food on an approximately 12:12 L:D cycle. For imaging experiments, *Drosophila* were genetically modified using the GAL4-UAS system to drive expression of the GCaMP6m calcium indicator in all glutamatergic neurons, including motor neurons, through OK371-GAL4 [31], [32], [33], [88]. All imaging experiments that formed the basis of the sequence analysis used 3^rd^ instar larvae in which OK371-GAL4 was recombined with GCaMP6m (OK371-GCaMP6m). All animals were reared at 21-24°C.

Individual 3^rd^ instar larvae were positioned and pinned on Sylgard-lined petri dishes dorsally through the mouthparts and posterior abdomen. Using fine scissors, an incision along the dorsal surface of the body wall enabled removal of all internal organs. The body wall was pinned flat exposing the CNS. The brain, suboesophageal ganglion (SOG), and ventral nerve cord (VNC) were dissected and pinned using fine tungsten wire (California Fine Wire, Grover Beach, CA) with remaining tissue removed before imaging. The dish was washed five times with Baines External Saline (BES) . Before imaging, fresh BES was applied containing (in mM): 135 NaCl, 5 KCL, 2CaCl_2_, 4 MgCl_2_, 5 TES and 36 Sucrose, pH 7.15. All imaging recordings were made within 15 min after excising the CNS.

Live preparations were imaged using an Olympus UPlanFL 10x INFINITY-corrected dipping lens with 0.3NA (UPLFLN10X2). Images were captured using WinFluor (version 4.1.9, University of Strathclyde, Strathclyde, UK) at 10fps. During imaging, all preparations were superfused with BES for up to 45 min. Images were stabilised against distal shifts using a Template Matching plugin in FIJI [89], [90]. Fluorescence values were manually extracted by placing regions of interest (ROIs) in thoracic (T1-3) and abdominal (A1-A8) hemi-segments within FIJI as reported in [1], [31]. Time series of ROI fluorescence values were extracted and pre-processed using rolling-ball averaging to create percentage change in fluorescence from baseline values (ΔF/F). Fluorescence data was visualised using DataView (11.17.1) [91]. Peaks in segmental activity were determined using the built-in hill-valley analysis facility in DataView.

### Classifying Fictive Activity

We measured sequences of fictive events. Behavioural sequences consisted of discrete categorical states where each element represented a single identified fictive motor event. Fictive activity was classified based on detecting peaks in motor neuron calcium signals and definitions of patterns of intersegment peak activity. The motor dynamics within the isolated VNC preparation were classified using Hill’s Valley thresholding to determine peaks within motor activity in each abdominal and thoracic hemi-segment alongside established definitions for each fictive behaviour [1], [31], [33] (Figure 7). Succinctly, fictive forward waves and fictive backward waves were determined by the existence of sequential Hill’s Valley-determined peaks proceeding symmetrically across the midline from either posterior-to-anterior or anterior-to-posterior directions, respectively. Fictive headsweeps were determined by computing a difference trace (left – right) in the T3 segment and determining anterior asymmetric activities as peaks or troughs that exceeded ±1% absolute threshold difference. Turning behaviours were defined as an anterior asymmetric activity immediately followed by a fictive backward event. Finally, anterior symmetric (i.e., anterior burst) or posterior symmetric (i.e., posterior burst) were determined by simultaneous peak detection of the left and right trace in T3-A2 and A6-8 respectively. Some anterior burst and posterior burst fictive activities are in varying numbers of anterior and posterior segments thus any peak detection in both the left and right trace within 0.5s of each other was determined as symmetric activities of each definition. In the calcium dataset, Hill’s Valley peak detection was performed after baseline smoothing within DataView [1]. A Python-based sorting algorithm (available in data repository) was used to isolate the peak times from DataView and to sort into temporal order. Ambiguous cases (e.g., lack of a segment peak, edge-cases) were flagged and manually reviewed. Manual corrections – deletions or additions of peaks – were performed, when necessary, in line with the fictive definitions highlighted above.

**Figure 7:**
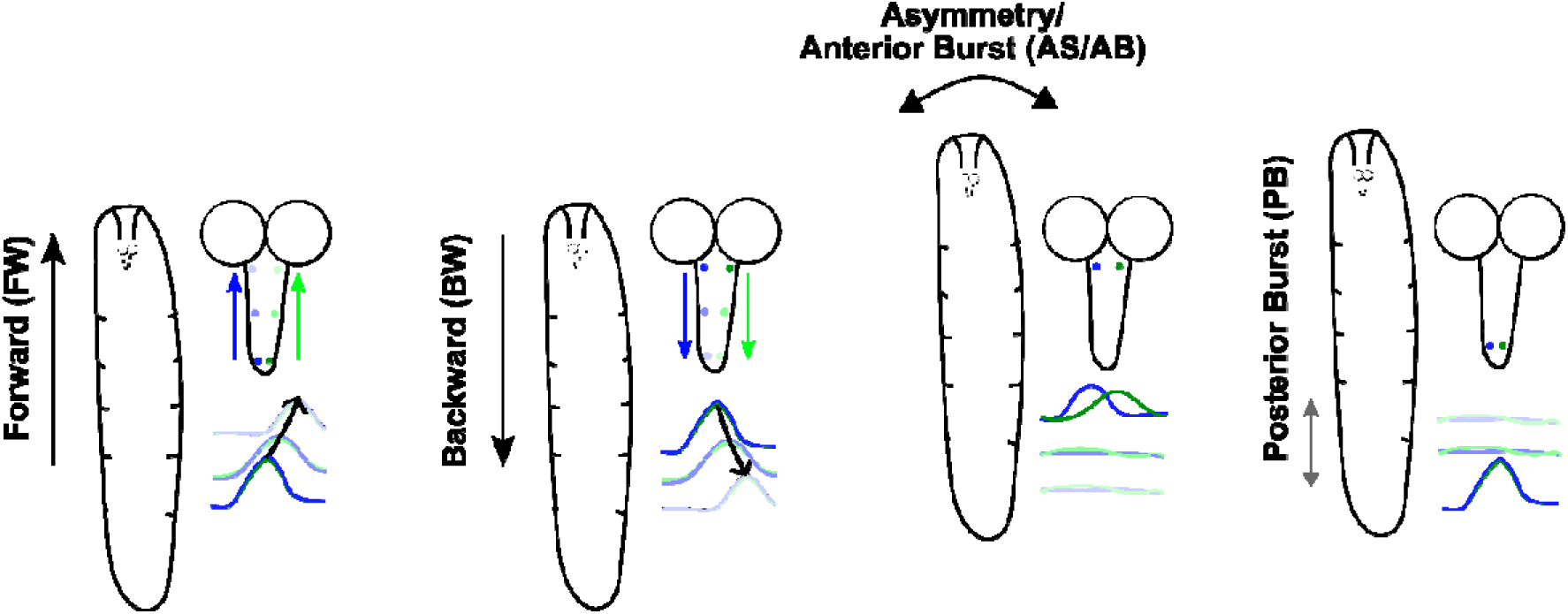
Fictive Behavioural Categories. The intact behaviour of the *Drosophila* larva is classified by different motor neuron activity sequences within the isolated ventral nerve cord (as originally noted in [31]). All subsequent calcium imaging analysis categorises sequential motor activity in either of the six fictive patterns: fictive forward (FW), fictive backward (BW), anterior asymmetric (AS), anterior bursting (AB), or posterior bursting (PB) events.

### Sequence Motif Analysis

#### Transition & Correlation Analysis

For each imaged preparation (), the scored fictive behaviours were converted into an ordered symbolic sequence. Pairwise transition counts () were calculated between each state () and its immediate subsequent state. Counts were row-normalised to generate a first-order transition probability matrix whereby each element represented the probability of transition between one state to another () (1). Transition matrices were computed independently for each preparation and then averaged across preparation or population-level visualisation and analysis. To assess whether the occurrence of different fictive behaviours covaried across preparations, we calculated the frequency of each fictive state within each preparation with pairwise Pearson correlations computed between behavioural frequency vectors across preparations. Correlation strength was reported, alongside associated Benjamini-Hochberg FDR-corrected p-values, and multiple comparisons were corrected using false discovery rate correction where appropriate using Python’s scipy.stats and statsmodels.stats.multitest.

Several transition and graph metrics were computed to explore dominant transition patterns and hub-like dynamics. Transition metrics were computed directly from transition probability distributions. Dominant transitions were defined as the largest transition probability from one state to its dominant successor. Maximum ongoing transition probability was used to measure if one state had one strongly preferred outgoing relationship. Entropy was used to measure the diversity of transition sources or destinations, with higher entropy indicating a more distributed transition profile. Transition entropy from one state (i) to another (*H*_i_) was calculated using the Shannon entropy formulation (2). High outgoing entropy indicates that a state branches into a more diverse set of successor states whereas high incoming entropy indicates that a state can be reached from a more diverse set of preceding states. Network analysis was done using Python’s *networkx* package.

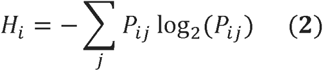

### Graph Node Metrics

To identify hub-like fictive states, selected graph metrics (*M*) were z-scored across states and averaged to produce a composite hub score (3). The composite score included weighted in-strength, weighted out-strength, total strength, outgoing entropy, PageRank, and betweenness centrality. Higher composite hub scores therefore indicate states that combine strong incoming and outgoing connectivity, diverse onward transitions, and centrality within the directed transition network. Specifically, weighted-in and weighing-out strength captures the total transition probability of entering or leaving a fictive state, excluding self-transitions. Weighted total strength measures the overall transition connectivity. Weighted in-degree and out-degree capture the number of non-zero incoming and outgoing transition links, respectively. PageRank estimated the importance of a node based upon incoming links from other important nodes [92]. A high PageRank state is one where the state itself is important in the transition graph. Eigenvector centrality was used to assess nodes which are highly connected to other highly connected nodes with Python’s *networkx* computing centrality based on centrality of predecessor nodes [93]. Succinctly, betweenness centrality measures the fraction shortest pathways between other node pairs that pass through a given note whereas closeness centrality measures how close a node is to all other nodes in a network.

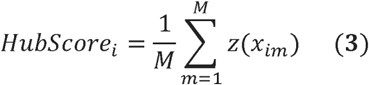

### N-Gram, Simulated Markov Datasets, & Motif Deviation Analysis

Multi-step fictive motifs were quantified using sliding-window n-gram counts range from 2-12 using Python’s *nltk* package. For each motif length, all contiguous fictive behavioural sequences were counted within each preparation and then summarised across preparations. The explicability of the one-step transition rules was evaluated through the construction of several distinct Markov order motifs against a unigram model, purely based on fictive behavioural frequencies. Expected motif frequencies were estimated under a first-order Markov model (4) in which each state depended only on the immediately preceding state and a variable-order Markov model where the next state (*s_n_*) depends upon longer histories (maximum of 3 prior states (*h*), 0.5 smoothing parameter). When a high-order context was insufficiently sampled, the additive/Laplace-smoothed (*a*) variable-order Markov model (5) backed off to a shorter context or to the unigram distribution. Transition probabilities were smoothed to avoid zero-probability events.

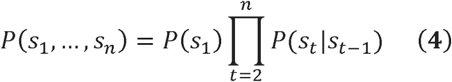

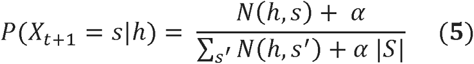

Simulated datasets were generated from the fitted transition model while preserving the empirical distribution of sequence lengths. Motif counts from simulated datasets were then used to estimate expected motif occurrence under the null model. Prediction accuracy was quantified by comparing empirical motif counts with model-predicted counts. For each motif length, relative prediction error and mean absolute percentage error were calculated across motifs alongside Pearson correlation being calculated between empirical motif counts and model-predicted motif counts to evaluate how well transition models preserved the relative abundance of motifs. The predictive performance between the three deviation models – unigram, 1°-order, and variable-order Markov model – was compared using held-out cross-entropy, calculated as the mean negative log-probability assigned to observed transitions in test preparations. Lower cross-entropy indicates better predictive performance. Further, model performance was evaluated using preparation-level cross-validation. For each fold, models were trained on a subset of preparations (20%) and evaluated on held-out preparations, preserving independence between training and test sequences. Cross-entropy values were compared across models using a Friedman repeated-measures test, followed by paired Wilcoxon signed-rank tests with Holm correction.

Motif enrichment was quantified by comparing empirical motif counts with null-model motif count distributions. Z-scores were calculated as the difference between observed and expected motif counts divided by the standard deviation of the null distribution. A shuffle-within-preparation null model was used to test whether motif enrichment could be explained by preparation-specific fictive behavioural frequencies alone thereby preserving the number and identity of fictive behaviours within each preparation but disrupting their temporal order. A first-order Markov null model was used to test whether motif enrichment could be explained by pairwise transition probabilities alone. Motifs that remained enriched or suppressed relative to this null were reported as evidence of higher-order sequence structure. Motif-level significance was assessed after false discovery rate correction. To avoid unstable enrichment estimates, motifs with very low expected counts, very low null variance, or extreme Z-scores caused by degenerate null distributions were excluded from enrichment visualisation. Significant motifs were further classified according to their fictive sequence structure, including repeated-state, return-to-state, oscillatory, and wave–distal switching motifs, to ease in identifying the types of higher-order trajectories enriched in the fictive transition landscape.

To test whether enriched motifs were simply caused by differences between recorded preparations, we repeated the motif-enrichment analysis using a preparation-specific first-order Markov null. From the observed fictive behavioural sequence, each preparation’s own transition matrix was used to generate a same-length simulated fictive sequence. Empirical motif counts were compared against the simulated motif-count distributions to identify enriched or suppressed motifs, with p-values corrected for multiple comparisons. Motifs that remained significant under this control were interpreted as higher-order sequence structures that could not be explained by each preparation’s own one-step transition rules.

### Stationarity Analysis

Stationarity analyses were performed within preparation-level sequences of temporally-ordered fictive behavioural events predominately using the Python package *ruptures* [94]. To assess coarse temporal stability, each preparation was divided into first and second halves of the sequence period. First-order transition matrices were computed separately for each half, and divergence between halves was quantified using Jensen–Shannon divergence (6). Jensen-Shannon distance was reported. To examine gradual changes in transition structure, transition matrices were estimated in sliding windows (40-event window, advancing by 20 events each time; *ruptures.Pelt(model=“rbf”)*, *pen=5*) across each preparation. Divergence between windowed transition matrices was used to quantify local drift in fictive transition dynamics. Given Jensen-Shannon (JS) divergences symmetrically measure the dissimilarity between probability distributions, this metric was used to compare transition probability distributions. The JS divergence was computed row-wise from the probability transition matric within each preparation’s halved-recording period. To determine whether apparent drift was driven by sequence length, we tested the relationship between each preparation’s drift magnitude and the number of scored events/transitions. Abrupt transition reorganisations were identified by searching each sequence for categorical points where behavioural frequency distributions before and after the candidate point diverged. Candidate change-points were defined by their local Jensen–Shannon divergence. Preparations were classified as stationary when median sliding-window JS ≤0.25 and split-half JS≤0.35. or strong drift when median sliding-window JS≥0.40 or split-half JS≥0.55 otherwise they were classified as moderate drift. Change-points were thus detected independently of the JS threshold using PELT (pen=5) [95]. JS divergence was then used post hoc to quantify the magnitude of local behavioural reorganisation around each detected change-point. The relationship between change-point timing and local divergence magnitude was assessed by correlation analysis. Local change-point divergences were compared within each preparation’s overall sliding-window drift score to assess if abrupt reorganisations within the fictive landscape reflected the same process as gradual drift. To assess whether non-stationarity was associated with changes in fictive behavioural composition, the proportion of each fictive state was calculated in sliding windows and averaged across preparations with uncertainty reported in SEM.

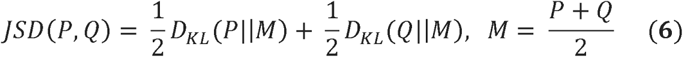

### Markov Memory Models

We compared 6 multinomial logistic-regression transition models (7), each predicting the next fictive state from a different set of historical predictors. Model 1 included only the current fictive state and therefore represented a first-order Markov model. Model 2 added the previous state to test whether two-step local context improved prediction. Model 3 added current run length, defined as the number of consecutive events for which the current state had persisted, to test whether local bout persistence shaped transition probability. Model 4 added state-specific since-last-state predictors, encoding the number of intervening fictive events since each state was last expressed, to test for categorical recency memory. Model 5 added recent-window summary predictors describing the diversity, balance, or composition of recent fictive activity. Model 6 combined all candidate memory predictors and represented the full-memory model. By comparing these models under the same log-loss, held-out evaluation framework, we assessed which form of fictive behavioural history best explained fictive transition structure. For simulation, each model was fitted to the full transition-level dataset using multinomial logistic regression. Categorical predictors were one-hot encoded and continuous predictors were standardised before model fitting (5000 maximum iterations with regularisation parameter of 1.0 and Lbfgs solver [96]). Model simulations were generated in closed loop. For each preparation, the simulated sequence was initialised with the empirical first state and then iteratively extended by sampling the next state from the model-predicted probability distribution. History-dependent predictors were recomputed from the simulated sequence at each step, and simulated sequences were matched to the empirical sequence lengths. 100 simulated datasets were generated per model. During closed-loop simulation, all history-dependent predictors were recalculated from the simulated sequence history. Since-last-state predictors encoded the event lag since each state was last expressed, whereas recent-window predictors summarised the composition of the most recent behavioural events. Empirical motif structure was quantified by counting contiguous 3-gram and 4-gram motifs across preparation-level sequences. Motifs with fewer than five empirical occurrences were excluded from motif-reproduction comparisons. For each simulated dataset, motif counts were calculated using the same procedure as for empirical sequences. Model motif counts were then averaged across simulation replicates. For each empirical and simulated dataset, motif enrichment was quantified relative to a first-order Markov expectation. The expected count of each motif was estimated from the frequency of its prefix and the 1°-order probability of its final transition. Motif enrichment was expressed as a Z-score, with positive values indicating enrichment and negative values indicating suppression relative to 1^st^-order expectations. Such analysis used an analytical first-order Markov expectation to provide a consistent motif-deviation score across empirical and model-generated sequences. Motif direction was assigned from the sign of the enrichment Z-score. Motifs with positive Z-scores were classified as enriched, and motifs with negative Z-scores were classified as suppressed. No strict threshold was enforced.

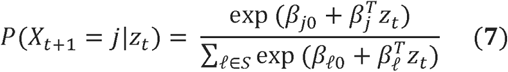

To ease in comprehension, multi-step fictive motifs were classified into arbitrary classes. Specifically, motifs were annotated by structural class to determine whether model performance was concentrated in recurrence-like motifs. Recurrence-like motifs included A→B→A patterns, A→B→A→B alternations, non-adjacent state recurrence, and motifs in which the first and last states matched. Recurrence-like motifs were defined as motifs in which a state reappeared after intervening events, including ABA and ABAB structures. Immediate self-persistence motifs were annotated separately. For each motif, we compared empirical motif count and enrichment Z-score with the corresponding model-simulated mean across replicate simulations. Model reproduction was summarised using the correlation between empirical and model motif Z-scores, direction-agreement fraction, and absolute motif-deviation error. Such metrics were calculated overall, separately by motif order, by enrichment direction, by motif class, and for recurrence-like motifs. To test whether models captured both motif enrichment and suppression, empirical and model direction labels were compared using direction-agreement summaries and enriched/suppressed confusion matrices. To visualise motif-specific reproduction, we selected the strongest empirically enriched and suppressed recurrence-like motifs and compared their empirical enrichment scores with model-simulated scores from each model.

### Hidden State Markov Model

Fictive behavioural sequences were represented as ordered categorical observations, with one sequence per preparation. Observations were mapped onto a common behavioural alphabet comprising forward waves (FW), backward waves (BW), anterior bursts (AB), posterior bursts (PB), and asymmetric activity (AS). End labels and unclassified events were excluded before modelling. To infer latent network regimes underlying observed fictive behaviours, we first fitted categorical hidden Markov models (HMMs), in which each hidden state emitted observable fictive behaviours with state-specific multinomial emission probabilities. Within Python, we used the *hmmlearn* package’s *hmm.CategoricalHMM*.

Candidate HMMs containing 2–9 hidden states were fitted to the preparation-level sequences. Model dimensionality was evaluated using Akaike information criterion (AIC), Bayesian information criterion (BIC), and preparation-level cross-validated log-likelihood. Cross-validation was performed at the preparation level so that all events from a given preparation were assigned either to the training set or to the held-out set, avoiding train–test leakage between adjacent events from the same recording. For each candidate state number, models were fitted using expectation–maximisation for a maximum of 100 iterations, and the best-fitting solution was selected from repeated random initialisations by maximising training log-likelihood. The selected HMM contained six hidden states, which were subsequently interpreted from their emission probabilities, occupancy, transition probabilities, and decoded Viterbi paths.

To determine whether the standard HMM memoryless-dwell assumption was appropriate, we examined state-specific dwell distributions. For each preparation, consecutive runs of identical Viterbi-decoded hidden states were collapsed into dwell segments, and dwell length was defined as the number of consecutive observed fictive events assigned to the same hidden state before transition to another hidden state. Under a standard discrete-time HMM, dwell durations are geometrically distributed because the probability of leaving a state is constant at each step. We therefore compared each empirical state-specific dwell distribution against the geometric dwell distribution implied by the fitted HMM self-transition probability. Specifically, for each state (,), the HMM-implied probability of leaving the state was defined as *q_Z_*, where *A_ZZ_* is the fitted self-transition probability. The expected dwell distribution was therefore 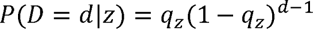, where (*d*) is dwell length in events.

State-specific deviations from geometric dwell structure were assessed using three criteria: visual inspection of the empirical and fitted dwell probability mass functions, comparison of empirical and geometric discrete hazard functions, and BIC comparison between a single geometric dwell model and a two-component dwell model. The discrete hazard was calculated as 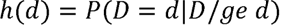, such that a constant hazard indicates a memoryless geometric dwell process. A hidden state was considered non-geometric if it showed a non-constant hazard and if the two-component dwell model improved BIC relative to the single geometric model. This analysis identified one broad-dwell AS-associated state, denoted ε, whose dwell distribution was poorly described by a single geometric process and instead contained both short-dwell and long-dwell components.

To represent this non-geometric persistence explicitly, we converted the fitted HMM into a hidden semi-Markov model (HSMM). In the HSMM, latent states retained categorical emission probabilities, but dwell durations were sampled explicitly from state-specific empirical dwell distributions rather than being imposed by a geometric self-transition process. For states with approximately geometric dwell structure, the HSMM used the empirical state-specific dwell probability mass function estimated from Viterbi-decoded dwell segments. For the non-geometric ε state, dwell segments were split into short- and long-dwell substates using the minimum between the short-dwell peak and long-dwell tail of the empirical dwell distribution. Operationally, ε dwell segments of one or two consecutive events were assigned to the short-dwell substate ε(s), whereas ε dwell segments of three or more consecutive events were assigned to the long-dwell substate ε(l). This threshold was chosen before posterior predictive simulation, based only on the empirical ε dwell distribution, and was not optimised to improve motif reproduction or downstream transition statistics.

After splitting ε into ε(s) and ε(l), each ε-associated dwell segment inherited the original ε emission identity but was assigned to ε(s) or ε(l) according to its dwell duration. Emission probabilities for ε(s) and ε(l) were then re-estimated from the observations assigned to each substate. The transition matrix was also re-estimated from the resulting substate sequence after collapsing consecutive dwell segments, so that transitions represented movement between latent regimes rather than repeated occupancy within the same dwell. Self-transitions were not used to generate persistence in the HSMM; instead, persistence was generated by the explicit dwell-duration distribution assigned to each state. Thus, the HSMM separated two sources of latent-state organisation: the probability of entering a state and the probability of remaining in that state for a given dwell duration.

HSMM simulations were generated by sampling an initial latent state from the empirical initial-state distribution, sampling a dwell duration from that state’s fitted dwell distribution, emitting observations from the state-specific multinomial emission distribution for the sampled dwell, and then sampling the next latent state from the fitted between-state transition matrix. This process was repeated until the simulated sequence reached the empirical preparation length, after which the sequence was truncated to match that preparation exactly. Simulations were therefore matched to the empirical distribution of preparation lengths while allowing latent-state persistence to arise from fitted dwell distributions rather than geometric self-transition probabilities. Posterior predictive checks were performed using 300 HSMM-simulated datasets. For each simulated dataset, we recalculated state occupancy, dwell statistics, event-level and bout-collapsed transition probabilities, transition entropy, motif counts, motif-enrichment scores, and temporal drift statistics. Empirical values were compared against the corresponding simulated posterior predictive distributions using 95% simulation intervals and posterior predictive p-values, with Benjamini–Hochberg false-discovery-rate correction applied across scalar posterior predictive checks.

### Sustainability Statement

Using the carbon calculator WillCO_2_st [97], [98], we estimate that all direct experimental work presented here generated approximately 608.9g CO_2_e using electricity provided by the South Scotland UK National Grid. In addition, using CodeCarbon [99], we estimate that all computational analysis and statistical modelling work generated 1.05kg CO_2_e based on the average South Scotland UK National Grid of 0.05 kg CO_2_e per kWh.

## Data Availability

The research data underpinning this publication can be accessed at https://doi.org/10.17630/f32df353-5e58-420f-821c-dd1a3bf11e9f.

## Author’s Contribution

S.R.P. and W.V.S. conceived the study. W.V.S conducted all experiments, analysis, and visualisation. W.V.S. and S.R.P finalised the figures. S.R.P. supervised the project.

## Acknowledgements

This project was made possible by an Industrial CASE PhD studentship (UKRI Biotechnology and Biological Sciences Research Council (BBSRC) grant number BB/T00875X/1) to WVS.

## Supplementary Information

**Supplementary Table 1.** Mean and Standard Error Transition Probabilities (% of total) between Fictive. Events Mean transition probabilities based on average between each preparation’s mean transition probabilities with standard error for each transition probability shown. **Bold** denotes the FDR-dominant transition destination.

|  |  | To |  |  |  |  |
| --- | --- | --- | --- | --- | --- | --- |
|  |  | AB | AS | BW | FW | PB |
| From | AB | 8.7 ± 1.3 | 25.3 ± 2.2 | 12.4 ± 1.9 | 11.5 ± 1.8 | **42.1 ± 2.7** |
|  | AS | 4.8 ± 0.5 | 30.9 ± 1.8 | 18.4 ± 1.4 | 9.8 ± 1.5 | 36.1 ± 2.1 |
|  | BW | 5.5 ± 0.8 | **70.6 ± 2.6** | 17.9 ± 1.7 | 1.2 ± 0.7 | 4.8 ± 1.2 |
|  | FW | 4.9 ± 1.1 | **50.7 ± 3.3** | 2.7 ± 1.2 | 4.8 ± 1.1 | 36.8 ± 3.3 |
|  | PB | 18.3 ± 1.7 | **47.7 ± 2.4** | 13.0 ± 1.2 | 7.8 ± 1.1 | 13.1 ± 1.2 |

**Supplementary Table 2.** Significant Pairwise Transition Probabilities Between Fictive Events. All comparisons are non-parametric Friedman test and Wilcoxon signed-rank test comparing “from”-“to A” vs “from”-“to B”. All fictive behaviours are present: forward (FW), backward (BW), anterior burst (AB), anterior asymmetric (AS), and posterior burst (PB) fictive patterns. Only significant results are detailed. BH-FDR within each A | B are calculated. Exclusively p-values < .05 are denoted. Green box indicates the median transition probability from the fictive event to A is significantly greater than B. Red box indicates the median transition probability from the fictive event to A is significantly lower than B.

| From | To A | To B |  |  |  |  |
| --- | --- | --- | --- | --- | --- | --- |
|  |  | AB | AS | BW | FW | PB |
| AB | AB | - | .0000 | - | - | .0000 |
|  | AS |  | - | .0000 | .0000 | .0006 |
|  | BW |  |  | - | - | .0000 |
|  | FW |  |  |  | - | .0000 |
|  | PB |  |  |  |  | - |
| AS | AB | - | .0000 | .0000 | .0021 | .0000 |
|  | AS |  | - | .0000 | .0000 | - |
|  | BW |  |  | - | .0000 | .0000 |
|  | FW |  |  |  | - | .0000 |
|  | PB |  |  |  |  | - |
| BW | AB | - | .0000 | .0000 | .0000 | - |
|  | AS |  | - | .0000 | .0000 | .0000 |
|  | BW |  |  | - | .0000 | .0000 |
|  | FW |  |  |  | - | .0000 |
|  | PB |  |  |  |  | - |
| FW | AB | - | .0000 | .0193 | - | .0000 |
|  | AS |  | - | .0000 | .0000 | .0332 |
|  | BW |  |  | - | .0332 | .0000 |
|  | FW |  |  |  | - | .0000 |
|  | PB |  |  |  |  | - |
| PB | AB | - | .0000 | .0252 | .0000 | .0345 |
|  | AS |  | - | .0000 | .0000 | .0000 |
|  | BW |  |  | - | .0011 | - |
|  | FW |  |  |  | - | .0000 |
|  | PB |  |  |  |  | - |

**Supplementary Table 3.** Median entropy (H) and confidence intervals (CI) per start state from isolated preparations that exhibited >2 of the fictive pattern (N).

| From | Median H | Lower CI | Higher CI | N |
| --- | --- | --- | --- | --- |
| AB | 1.54 | 1.43 | 1.67 | 72 |
| AS | 1.78 | 1.74 | 1.83 | 86 |
| BW | 1.49 | 1.40 | 1.57 | 85 |
| FW | 1.30 | 1.27 | 1.37 | 83 |
| PB | 1.83 | 1.75 | 1.91 | 80 |

**Supplementary Figure 1.**
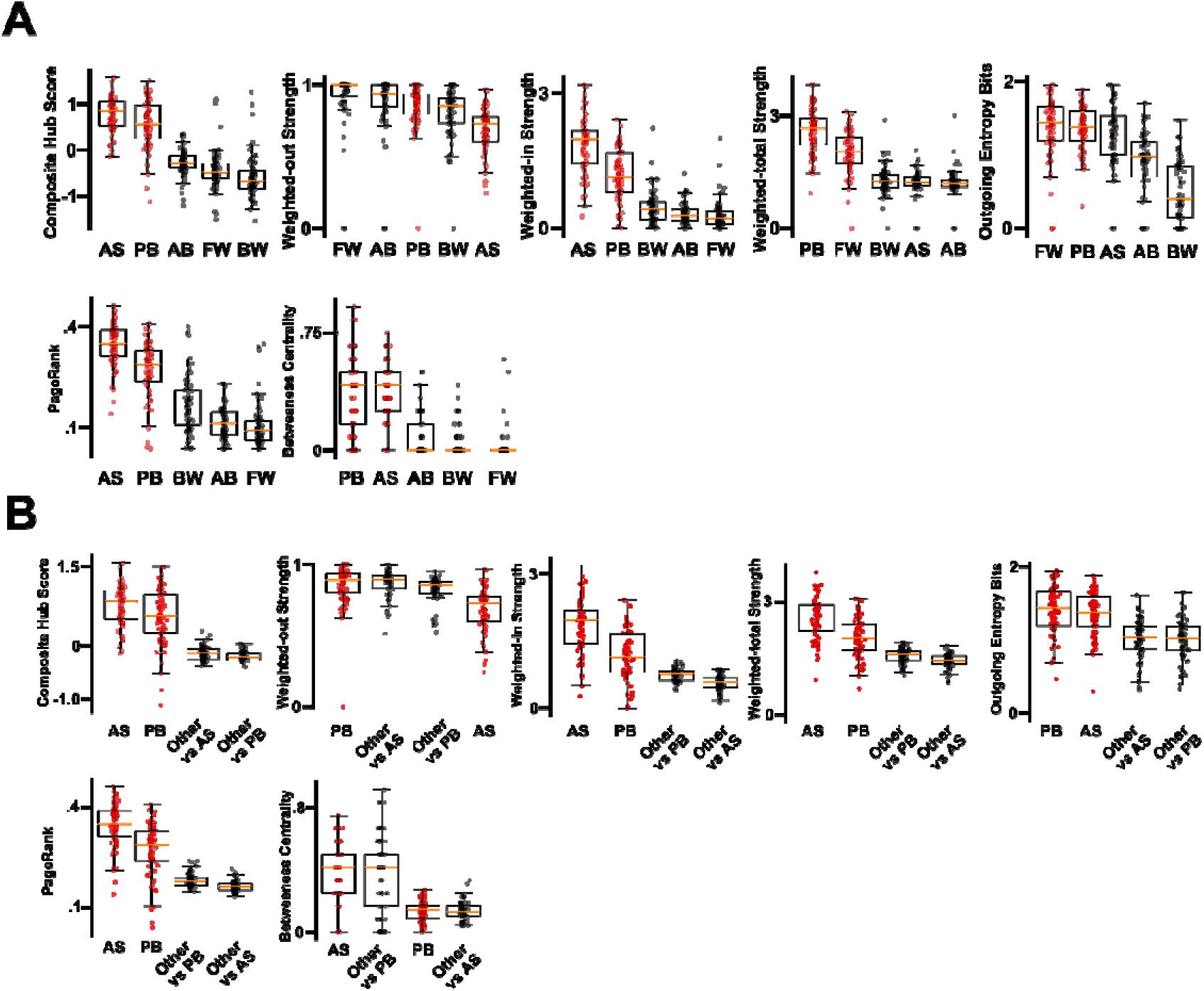
*Hub-liked Metrics Quantifying Fictive States* (**A**) Graph metrics (composite hub score, weighted out-strength, weighted-in strength, weighted total strength, outgoing entropy, Page Rank, betweenness centrality) across all fictive states (posterior burst (PB), anterior asymmetry (AS), forward wave (FW), backward wave (BW), and anterior burst (AB)). Across metrics, PB and AS are exhibit higher hub-associated values than FW, BW, or AB indicating that distal fictive activity states occupy a more central position in the transition landscape. (**B**) Metric value of anterior asymmetry (AS) and posterior burst (PB) against the mean value of non-hub states within the same preparation. Paired comparison shows AS and PB are generally elevated related to other non-hub states, thus AS/PB may form distal hub-like substates in the fictive motor transition network.

**Supplementary Figure 2.**
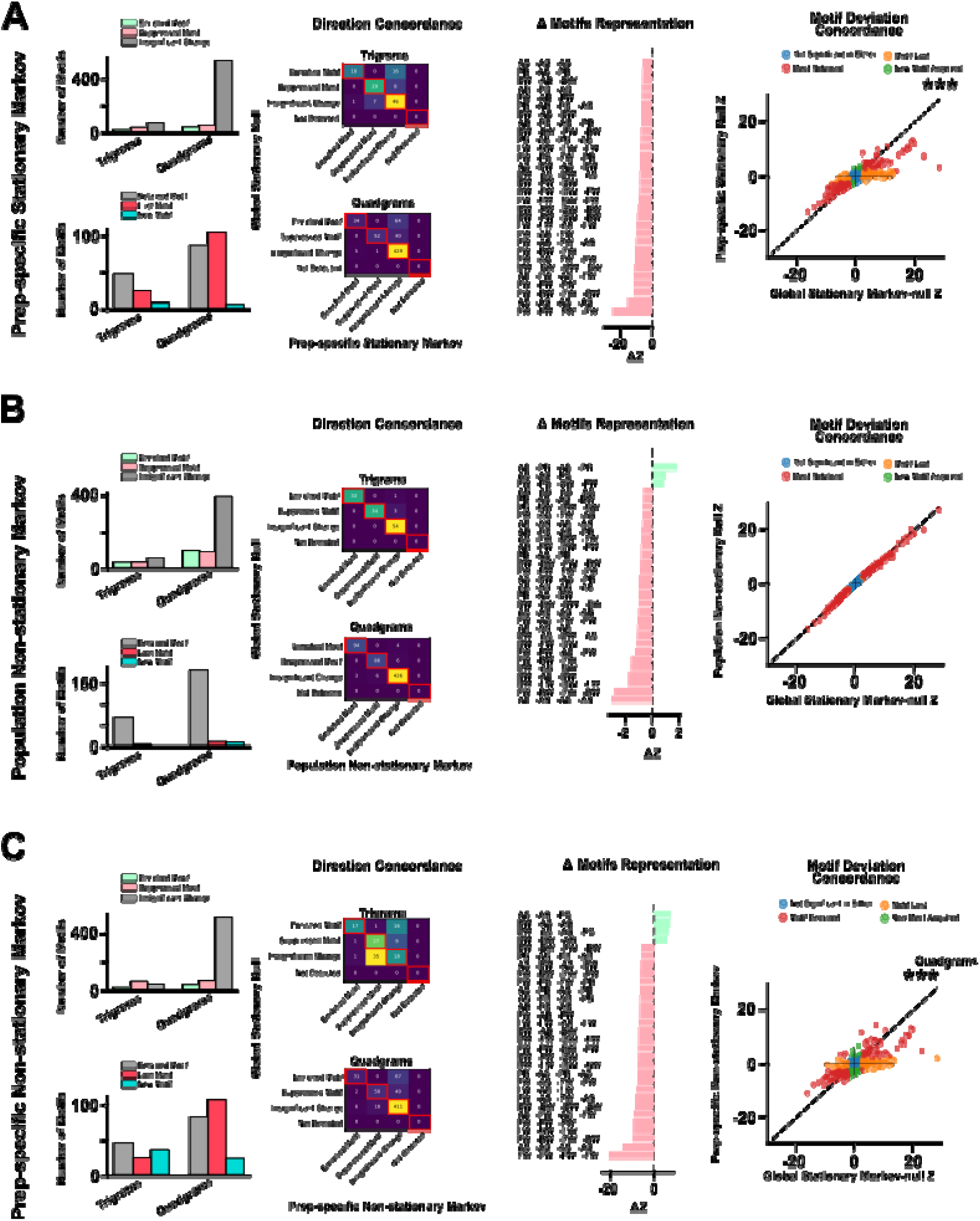
Non-stationarity Markov Null Preserves Multi-Step Fictive Motifs (**A**) Preparation-specific stationary Markov null comparisons against original Markov analysis. (**B**) Population-level non-stationary Markov null comparisons against original Markov analysis demonstrates even If the population transition matrix changes over time, multi-step motifs still deviate beyond a time-varying first-order Markov process. (**C**) Preparation-specific non-stationary Markov null comparisons against original Markov analysis demonstrates that even considering each preparation’s own time-varying one-step fictive transition riles, core multi-step motifs remain enriched or suppressed.

**Supplementary Figure 3.**
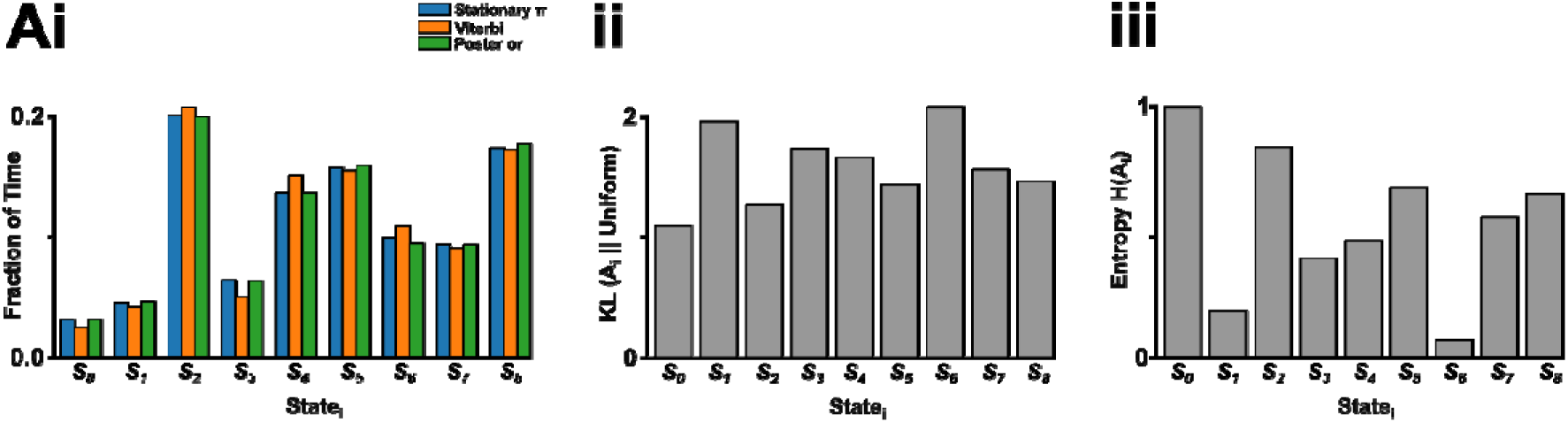
Comparable Alternative Base and Information Flow Metrics for Hidden-state Flow Diagrams (**A**) The hidden-state diagram for (**i**) probability flow, (**ii**) symmetric flow, (**iii**) surprise flow, (**iv**) mutual information flow, and (**v**) macrostate flow where edges >1% are displayed. Each substate is positioned relative to mean occupancy within the control fictive sequence data. Note the large similarity within the hidden-state diagram irrespective of elected metric. (**Bi**) The fraction of time each substate occupies showing non-uniform stationary distribution (*π*), Viterbi counts, and posterior counts thus emissions are not strongly forcing any particular path meaning transitions dominate occupancy with (Bii) high KL divergence from uniform (KL values > 1.5 for all substates), and (**Biii**) non-uniform entropy (0.14-1.07) indicative of mostly isolated “sticky” states.

**Supplementary Figure 4.**
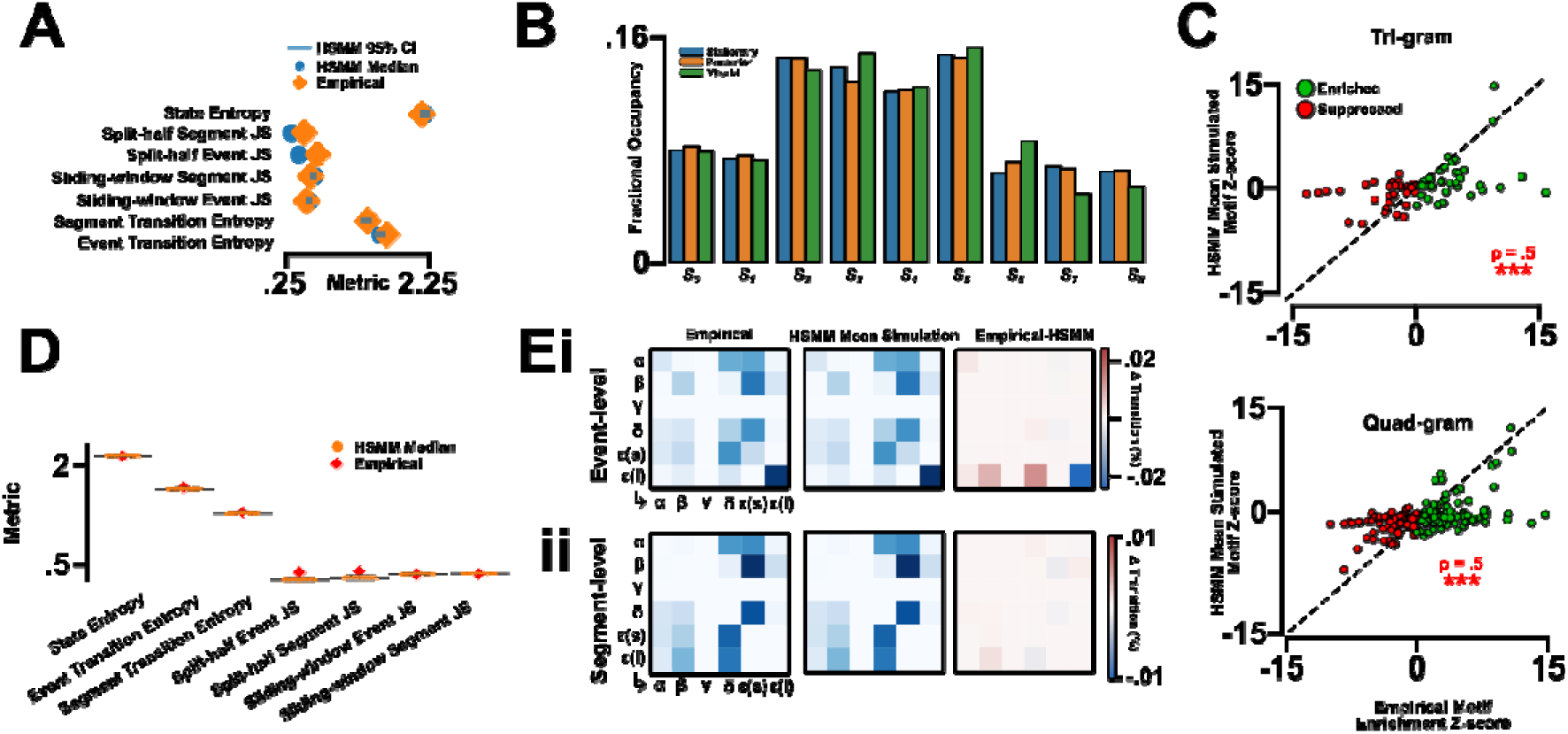
Posteriori & Robustness Checks for the HSMM (**A**) HSMM posterior predictive checks demonstrating strong continuity between empirical transition properties in fictive activity and the HSMM median. (**B**) HSMM hidden-state occupancy validation. (**C**) HSMM mean simulated motif z-score for tri- and quad-grams relative to empirical motif enrichment Z-score showing significant positive correlation between HSMM prediction and empirical mutli-step motif reality. (**D**) HSMM simulation stability and empirical fit of empirical transition properties in fictive activity. (**E**) Transition matrix between (**i**) event-level activity whereby each observed fictive behaviour is used for transition probability per-prep calculations and (**ii**) sequence-level activity whereby only transition between prolonged recurrence motif is examined during transition probability (e.g., FW → FW → FW → AB is catagorised under the FW → AB transition probability).

